# Telomeres control human telomerase (*hTERT*) expression through non-telomeric TRF2

**DOI:** 10.1101/2023.10.09.561466

**Authors:** Antara Sengupta, Soujanya Vinayagamurthy, Drishti Soni, Rajlekha Deb, Ananda Kishore Mukherjee, Subhajit Dutta, Jushta Jaiswal, Mukta Yadav, Shalu Sharma, Sulochana Bagri, Shuvra Shekhar Roy, Priya Poonia, Ankita Singh, Divya Khanna, Amit Kumar Bhatt, Akshay Sharma, Suman Saurav, Rajender K Motiani, Shantanu Chowdhury

## Abstract

The function of the human telomerase reverse transcriptase (*hTERT*) in the synthesis and maintenance of chromosome ends, or telomeres, is widely understood. Whether and how telomeres, on the other hand, influence *hTERT* regulation is relatively less studied. We found *hTERT* was transcriptionally altered depending on telomere length (TL). This resulted from TL-dependent binding of TRF2 between telomeres and the *hTERT* promoter. *hTERT* promoter-bound TRF2 was non-telomeric and did not involve the looping of telomeres to the *hTERT* promoter. Cell lines from different tissue types (fibrosarcoma (HT1080), colon cancer (HCT116), and breast cancer (MDA-MB-231), engineered for either telomere elongation/shortening gave increase/decrease in *hTERT*, respectively. Mechanistically, we show *hTERT* promoter-bound non-telomeric TRF2 recruits the canonical PRC2-complex inducing repressor histone H3K27-trimethylation in a TL-dependent fashion. This was further supported by TL-dependent promoter activity from an exogenously inserted *hTERT* reporter. Increase in TL over days followed by gradual decline, resulted in activation followed by repression of *hTERT* in a concerted manner, further implicating TL as a key factor for *hTERT* regulation. Notably on reprogramming primary fibroblasts to induced pluripotent stem cells (iPSCs), TRF2 loss from the *hTERT* promoter was evident along with telomere elongation and *hTERT* upregulation. Conversely, on telomere shortening in iPSCs, *hTERT* promoter-bound TRF2 was restored with marked reduction in *hTERT* further supporting the causal role of TL in *hTERT* transcription. Mechanisms of tight control of *hTERT* by TL shown here are likely to have major implications in telomere-related physiologies, particularly, cancer, ageing and pluripotency.

**Teaser:** Telomere length controls *hTERT* expression by modulating TRF2 distribution and PRC2-mediated repression, highlighting a self-regulatory mechanism in cancer.

## Introduction

Telomeres are nucleoprotein structures comprising guanine-rich DNA sequences at the end of linear chromosomes(1–3). The catalytic subunit of the ribonucleoprotein telomerase, <u>h</u>uman <u>te</u>lomerase reverse transcriptase (hTERT) is necessary for replicating telomeric repeats to maintain telomeres(4–7). Most cancer cells, in contrast to normal adult somatic cells, maintain telomeres through reactivation of *hTERT* (∼90% of cancers); alternative lengthening of telomeres (ALT) is used in other cases(8–13). *hTERT* therefore is tightly regulated at the epigenetic, transcriptional and translational levels(14,15).

The role of *hTERT* in the synthesis and maintenance of telomeres has been extensively studied(3,5,7,14,16,17). Relatively recent work suggests how telomeres, on the other hand, might affect *hTERT* regulation. Physical looping of telomeres reported as the <u>T</u>elomeric <u>P</u>osition <u>E</u>ffect-<u>o</u>ver-long-<u>d</u>istance (TPE-OLD) was shown to extend up to 10 Mb of telomeres (sub-telomeric). The resulting telomeric heterochromatinization gave TL-dependent repression of sub-telomeric genes. The *hTERT* loci(∼1.2 Mb away from telomeres) was shown to be repressed through (TPE-OLD)(18–20); loss of looping in short telomeres induced *hTERT*. Notably, another conceptually distinct mechanism of TL-dependent gene regulation was reported which influenced genes spread throughout the genome: expression of genes distal from telomeres (for instance 60 Mb from the nearest telomere) was altered in a TL-dependent way, but without physical telomere looping interactions. First, non-telomeric binding of TRF2 was found to be extensive (>20,000 sites), spread across the genome including promoters, and affected the epigenetic regulation of genes(21–23).Second, the shortening or elongation of telomeres led to the release or sequestration of telomeric TRF2, respectively, thereby increasing or decreasing the availability of TRF2 at non-telomeric promoters and affecting gene expression(21,24). The telomeric sequestration-dependent partitioning of telomeric versus non-telomeric TRF2 was proposed as the <u>T</u>elomere-<u>S</u>equestration-<u>P</u>artitioning (TSP) model, linking global gene regulation to telomeres (Fig 1) (21,25–27). Further, we found promoter-bound TRF2, not associated with telomeres (or non-telomeric), repressed *hTERT*(28). Based on these, here we tested the hypothesis non-telomeric TRF2 binding at the *hTERT* promoter was TL-dependent. If so, this would molecularly link telomeres to *hTERT* regulation.

**Fig 1.**
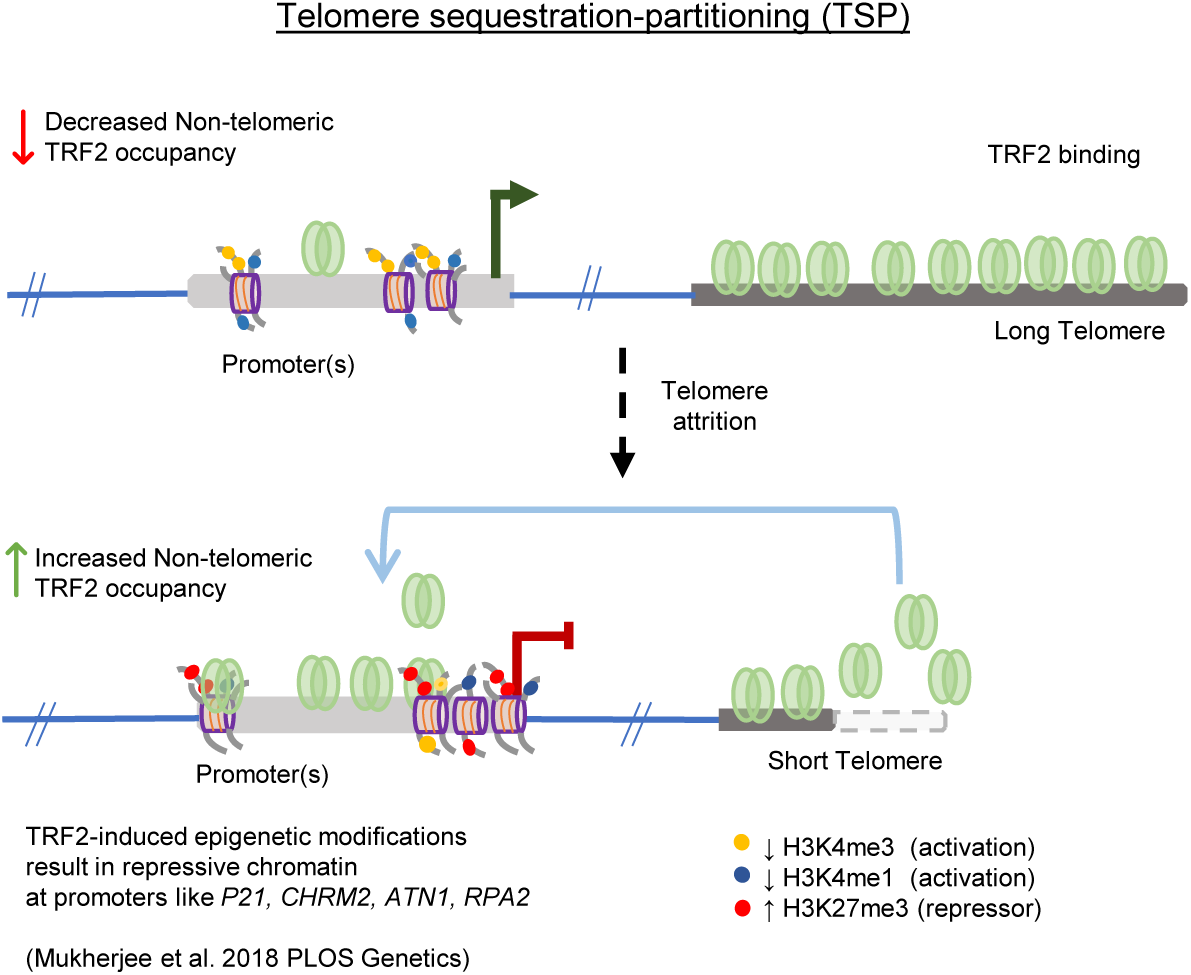
Graphical representation of the Telomere Sequestration Partitioning (TSP) model. The binding of TRF2 outside of telomeric regions (non-telomeric) is dependent on telomere length. When telomeres shorten, the occupancy of TRF2 at telomeres diminishes, resulting in increased TRF2 binding at non-telomeric sites. This shift in TRF2 distribution triggers epigenetic modifications at promoters showing telomere-dependent gene regulation.

Findings show TRF2 binding at the *hTERT* promoter increased with telomere shortening and downregulated *hTERT*. Conversely, reduced promoter TRF2 on telomere elongation upregulated *hTERT*; consistent with TSP described above. This was clear in multiple cell types engineered for either telomere elongation or shortening. An artificially inserted *hTERT* promoter (>40 Mb from telomere) showed TRF2-dependent and telomere-sensitive regulation, confirming the mechanism to be independent of physical telomere looping. For a temporal model, we elongated telomeres using doxycycline (dox) inducible cells and stopped elongation after a desired time. TRF2 binding at the *hTERT* promoter decreased as telomeres elongated over several days, and increased again with telomere shortening when dox treatment was stopped. Together, these show a heretofore unknown mechanism of telomere-dependent regulation of *hTERT*, the sole enzyme necessary for synthesizing telomeres, revealing a cellular feedback machinery connecting telomeres and *hTERT* expression.

## Results

### Telomere elongation upregulates *hTERT* through non-telomeric TRF2

To test if telomere length (TL) of cells affected TRF2 occupancy at the *hTERT* promoter, we first elongated telomeres in cancer cells under isogenic backgrounds (termed as <u>S</u>hort <u>T</u>elomere, ST or <u>L</u>ong <u>T</u>elomere, LT cells hereon). We used HT1080 fibrosarcoma cells (HT1080-ST) or the TL-elongated version HT1080-LT (with constitutive expression of *hTERT* and the RNA component *hTERC*; previously characterized by others and us, see Methods)(21,29). TL difference between HT1080-ST/LT was confirmed by flow cytometry (FACS, Fig 2A and qPCR based method, Supplementary Fig 1A, and telomerase activity in Supplementary Fig 1B). Chromatin-immunoprecipitation (ChIP) of TRF2 gave significantly reduced occupancy of TRF2 on the *hTERT* promoter in HT1080-LT compared to HT1080-ST cells (Fig 2B, ChIP-qPCR spanning 0-300 bp upstream of transcription start site (TSS)).

**Fig 2.**
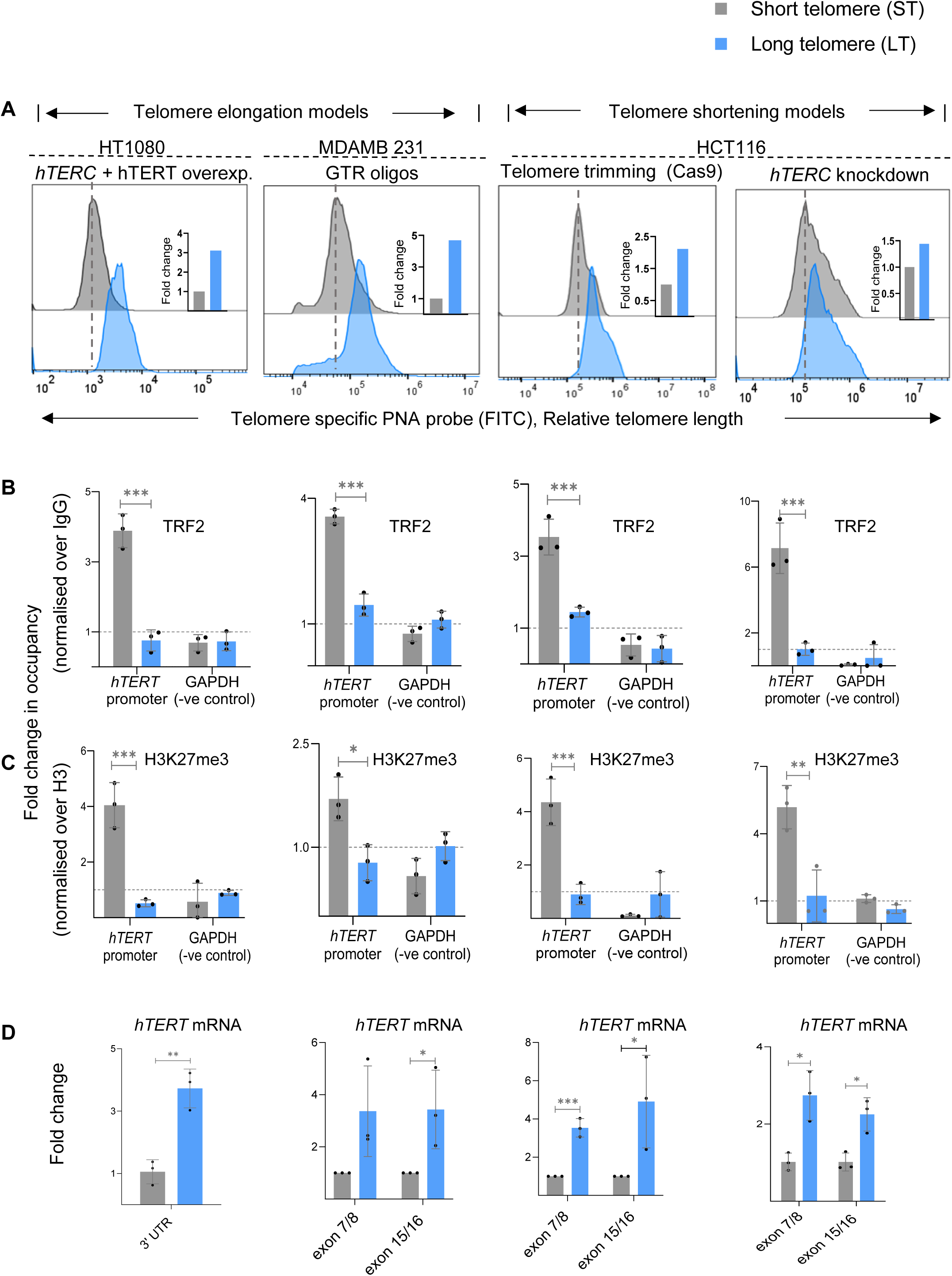
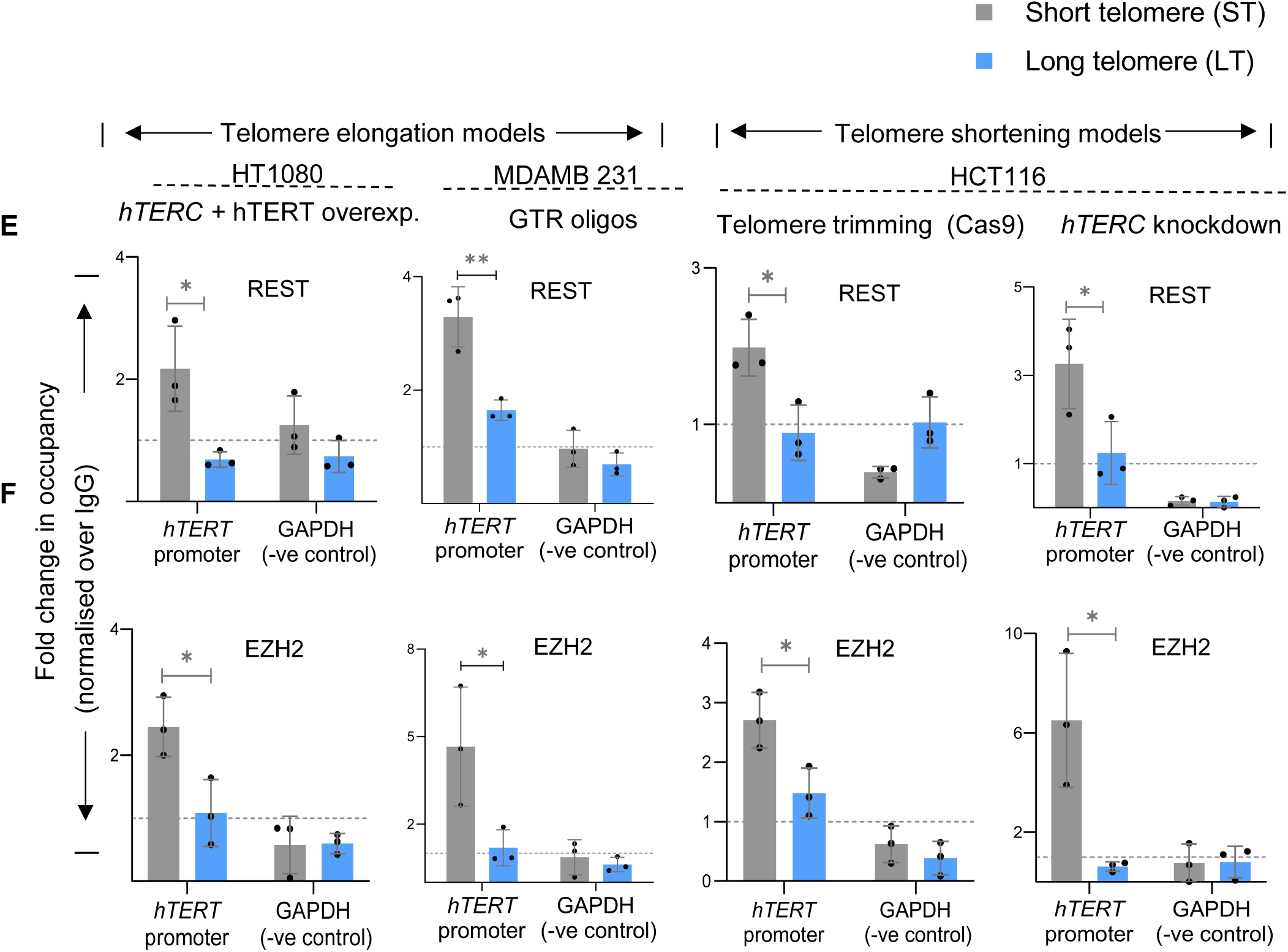
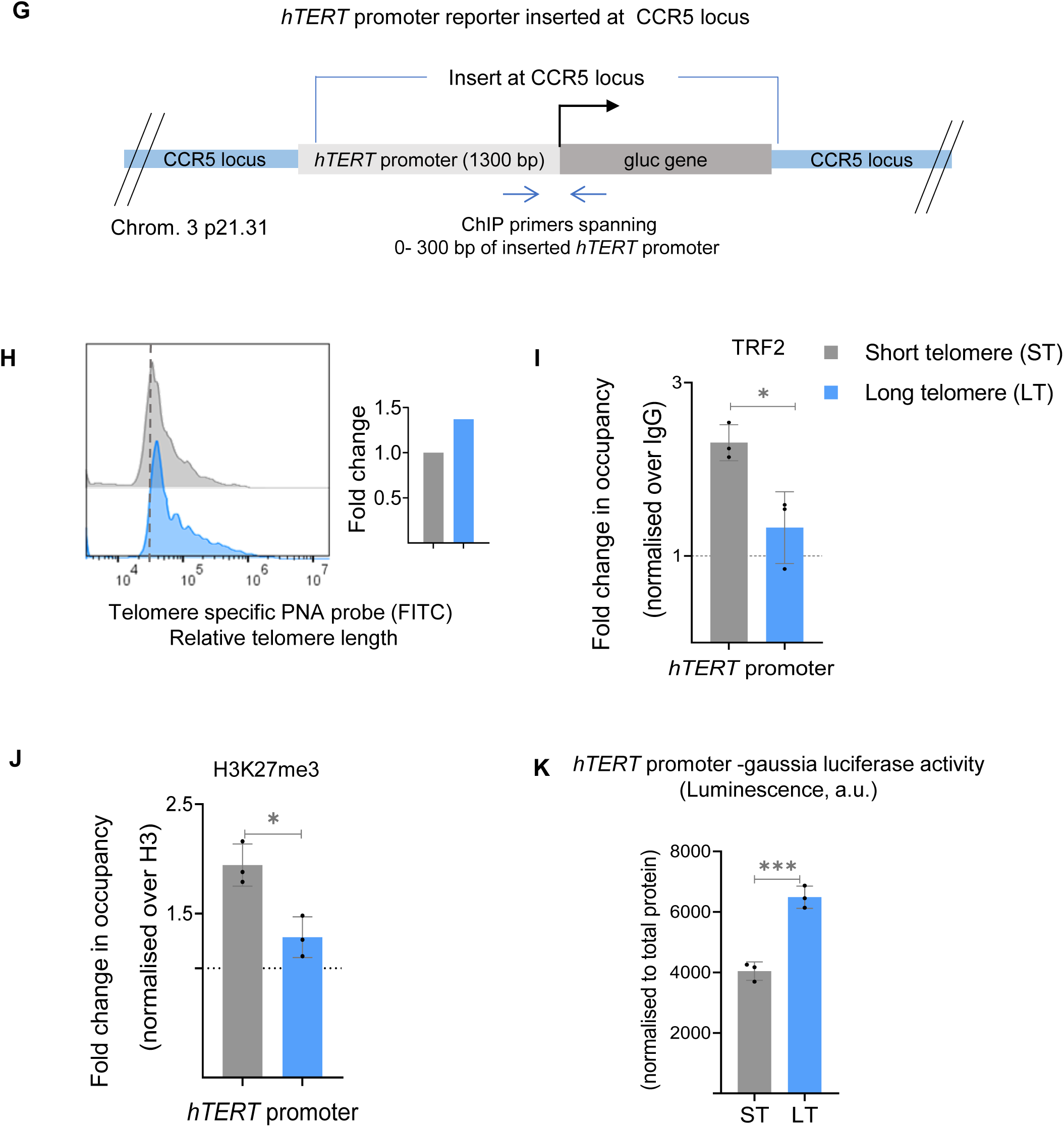
Telomere-dependent non-telomeric TRF2 binding at the *hTERT* promoter controls *hTERT* expression (A) Telomere length in isogenic cancer cell lines with short telomeres (ST, in grey) or long telomeres (LT, in blue) namely, HT1080-ST/LT, MDA-MB-231-ST/LT, and HCT116-ST/LT (Telomere trimming-Cas9 and *hTERC* knockdown) as determined by Flow cytometry (FACS). MDA-MB-231-ST/LT and HCT116-ST/LT Telomere trimming-Cas9 models were generated by telomerase-independent mode of TL alteration (See Methods). Relative fold change is shown in insets with FACS plots. (B-C) ChIP followed by qRT-PCR at the 0-300 bp *hTERT* promoter (upstream of TSS) for TRF2 (B) and H3K27me3 (C) in respective ST/LT cells as mentioned in (A); occupancy normalised to respective IgG or total Histone H3 (for H3K27me3). qPCR on the *GAPDH* promoter was used as the negative control in all cases. (D) *hTERT* mRNA expression by qRT-PCR in ST/LT cell line pairs as mentioned in (A), normalised to *GAPDH* or 18S mRNA levels. Primers specific to 3’UTR for endogenous *hTERT* were used for the HT1080-ST/LT system where telomerase was overexpressed for telomere elongation; primers for functional (reverse transcriptase domain) (exon7/8) and full length (exon 15/16) transcript were used for all other systems. MDA-MB-231-ST/LT and HCT116-ST/LT Telomere trimming-Cas9 models have been analysed as paired samples in each biological replicates. (E, F) ChIP followed by qRT-PCR at the *hTERT* promoter for REST (E) and EZH2 (F) in respective ST/LT cells as mentioned in (A); occupancy normalised to respective IgG. qPCR on the *GAPDH* promoter was used as the negative control in all cases. (G) Scheme depicting CRISPR modified HEK293T cells with 1300 bp *hTERT* promoter driving gaussia luciferase (Gluc) construct inserted at the CCR5 safe harbour locus. Scheme denotes ChIP primers used to study chromatin occupancy of 0-300 bp *hTERT* promoter region inserted at the exogenous locus. (H) Relative fold change in telomere length in *hTERT* promoter insert cells following telomere shortening determined by FACS; quantification in the right panel. (I, J) ChIP followed by qRT-PCR at the 0-300 bp *hTERT* promoter (upstream of TSS) insert at CCR5 locus for TRF2 (J), and H3K27me3 (K) in ST/LT cells. Occupancy normalised to respective IgG or total histone H3 (for H3K27me3). (K) *hTERT* promoter-gaussia luciferase activity in ST cells over LT cells from inserted exogenous with *hTERT* promoter. Reporter activity is presented as luminescence, (arbitrary units, a.u.) normalised to respective total protein levels. Error bars represent ± SDs from the mean of 3 independent biological replicates of each experiment. p values are calculated by unpaired t-test for all data except mRNA for MDA-MB-231-ST/LT and HCT116-ST/LT Telomere trimming-Cas9 models where paired t-test was performed. (*p < 0.05, **p < 0.01, ***p < 0.005, ****p < 0.0001).

TRF2-dependent H3K27 trimethylation(21), including at the *hTERT* promoter(28), was reported earlier by us: TRF2 silencing reduced the H3K27me3 mark at the *hTERT* promoter along with an increase in *hTERT* expression in the different cell lines tested(28). Here, we asked if TRF2-dependent *hTERT* regulation was TL-sensitive. With TL elongation, the H3K27me3 mark was significantly reduced in HT1080-LT relative to the ST cells (Fig 2C). As expected from reduced promoter H3K27 trimethylation, *hTERT* transcription was enhanced by >3 folds in HT1080-LT relative to ST cells; 3’UTR-specific primers were used to distinguish from exogenously expressed *hTERT* (Fig 2D).

To rule out any effect of exogenous *hTERT* (HT1080-LT cells above) we sought to elongate telomeres in an *hTERT-*independent way. Using a reported method, telomeres were elongated in MDA-MB-231 breast cancer cells with guanine-rich telomeric repeat (GTR) oligonucleotides (characterized by others and us, see Methods; termed MDA-MB-231-ST/LT; TL difference quantified using FACS, Fig 2A, and qPCR based method, Supplementary Fig 1A, and telomerase activity in Supplementary Fig 1B)(21,30). MDA-MB-231-LT cells had lower TRF2 binding at the *hTERT* promoter compared to MDA-MB-231-ST cells (Fig 2B). Accordingly, H3K27 trimethylation at the *hTERT* promoter was reduced and *hTERT* upregulated by ∼3-folds in MDA-MB-231-LT compared to MDA-MB-231-ST cells (Fig 2C-D). Together, TL-sensitive regulation of *hTERT* was consistent in both HT1080 and MDA-MB-231 cells on telomere elongation.

### Telomere shortening increased promoter TRF2 suppressing *hTERT* expression

As an alternative model, instead of elongation, we shortened TL in HCT116 colorectal carcinoma cells (with a higher average TL (∼5.6 kb) compared to HT1080 (∼4.5 kb) or MDA-MB-231 (∼3 kb) cells)(21,29,31). Two different methods were used for TL shortening:

(a) *hTERT*-independent, telomere-specific sgRNA guided CRISPR-Cas9 to trim telomeres (see Methods, scheme in Supplementary Fig 1C) or (b) depletion of the *hTERC* RNA (see Methods, TLs in HCT116-ST/LT quantified using FACS; Fig 2A, and qPCR based method, Supplementary Fig 1A, and telomerase activity in Supplementary Fig 1B). The ST cells in both TL-shortened models had higher TRF2 binding on the *hTERT* promoter compared to the HCT116-LT cells (Fig 2B). Also, we found a higher repressor H3K27me3 mark at the *hTERT* promoter, as expected from increased TRF2, in ST compared to its LT counterpart in both the HCT116 models (Fig 2C). As expected, an increase in the H3K27me3 repressor mark at the *hTERT* promoter resulted in >2-fold reduced *hTERT* expression in ST relative to the HCT116-LT cells (Fig 2D).

Previously, enhanced H3K27 trimethylation from TRF2-dependent binding of the RE1-silencing factor (REST) and EZH2 (the catalytic component of the PRC2 repressor complex) was reported at the *hTERT* promoter(28). Here we asked if the TL-sensitive H3K27 trimethylation resulted from altered REST/EZH2 binding at the *hTERT* promoter.

ChIP of REST showed enhanced occupancy at the *hTERT* promoter in ST relative to HT1080-LT cells (Fig 2E). EZH2 binding was also higher at the *hTERT* promoter in ST compared to HT1080-LT cells (Fig 2F). Similarly, in MDA-MB-231 cells, REST and EZH2 occupancy at the *hTERT* promoter was lower in LT compared to ST cells (Fig 2E-F). Further, an increase in REST and EZH2 binding at the hTERT promoter in ST relative to LT cells was clear in the HCT116-ST/LT cells (Fig 2E-F).

### Artificially inserted *hTERT* promoter is telomere sensitive

To further test TL-sensitive *hTERT* promoter activity we used a reporter cassette comprising 1300 bp of the *hTERT* promoter upstream of the *gaussia luciferase* (*Gluc*) gene. The reporter was inserted at the CCR5 safe-harbour locus, 46 Mb away from the nearest telomere, using CRISPR in HEK293T cells (Fig 2G). TL in HEK293T cells was shortened using telomere-specific sgRNA guided CRISPR-Cas9 to trim telomeres (Fig 2H, see Methods, scheme in Supplementary Fig 1C). Higher TRF2 binding at the inserted *hTERT* promoter in ST compared to the HEK293-LT cells was clear (ChIP-qRT primers specific to the CCR5-*hTERT* insert were used; Fig 2G, I). Consistent with this, H3K27me3 deposition was relatively high in ST cells (Fig 2J), and the *hTERT* promoter-driven Gluc reporter activity was reduced in the ST relative to the LT cells (Fig 2K). Together, these show TL-dependent *hTERT* regulation through non-telomeric TRF2 as a likely intrinsic mechanism.

### Temporally elongated or shortened telomeres up or downregulate *hTERT*

We next studied the effect of time-dependent (temporal) TL elongation/shortening, *i.e.*, TL increase followed by a decrease in a continuous way. For this, we made stable HT1080 cells with doxycycline-inducible hTERT. Following induction of hTERT and telomerase activity (Day 0), we checked TL through days 6/8/10/16 and 24 (Supplementary Fig 2A left panel). Gradual increase in TL to >5-fold by Day 10 (relative to Day 0) was evident; beyond Day 10 despite dox induction (and relatively high telomerase activity) further TL elongation was not seen (Supplementary Fig 2A-B left panels; ++/-- denotes presence/absence of dox, respectively).

For detailed analysis we focused on days 0,10 and 24 as representative time points; with discontinuation of dox after Day 10 (Dox-HT1080, Fig 3A). Following the withdrawal of dox after Day 10, TL returned to roughly within 1.6-fold of the initial (Day 0) state by Day 24 (Fig 3B and Supplementary Fig 2A left panel).

**Fig 3.**
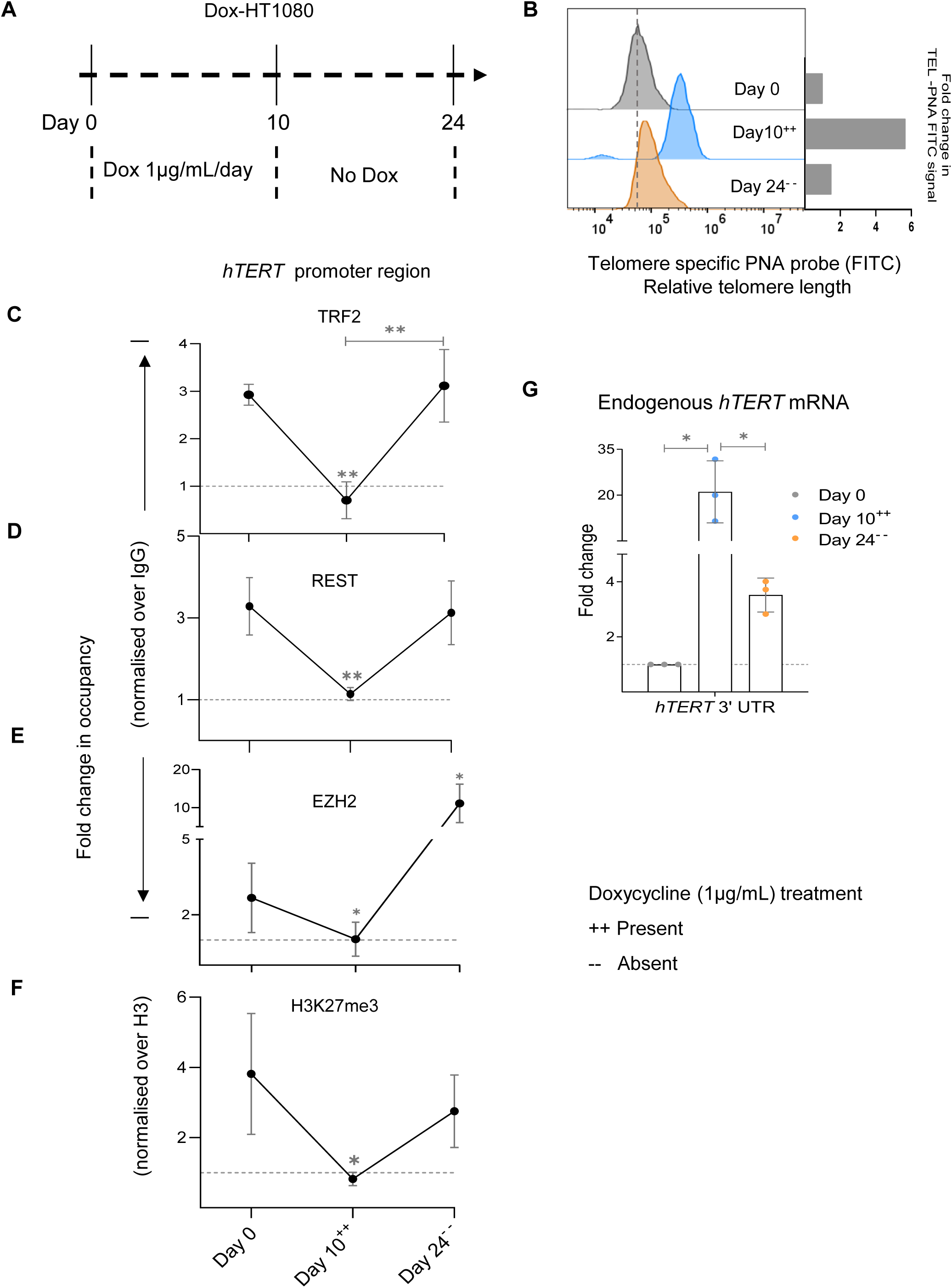
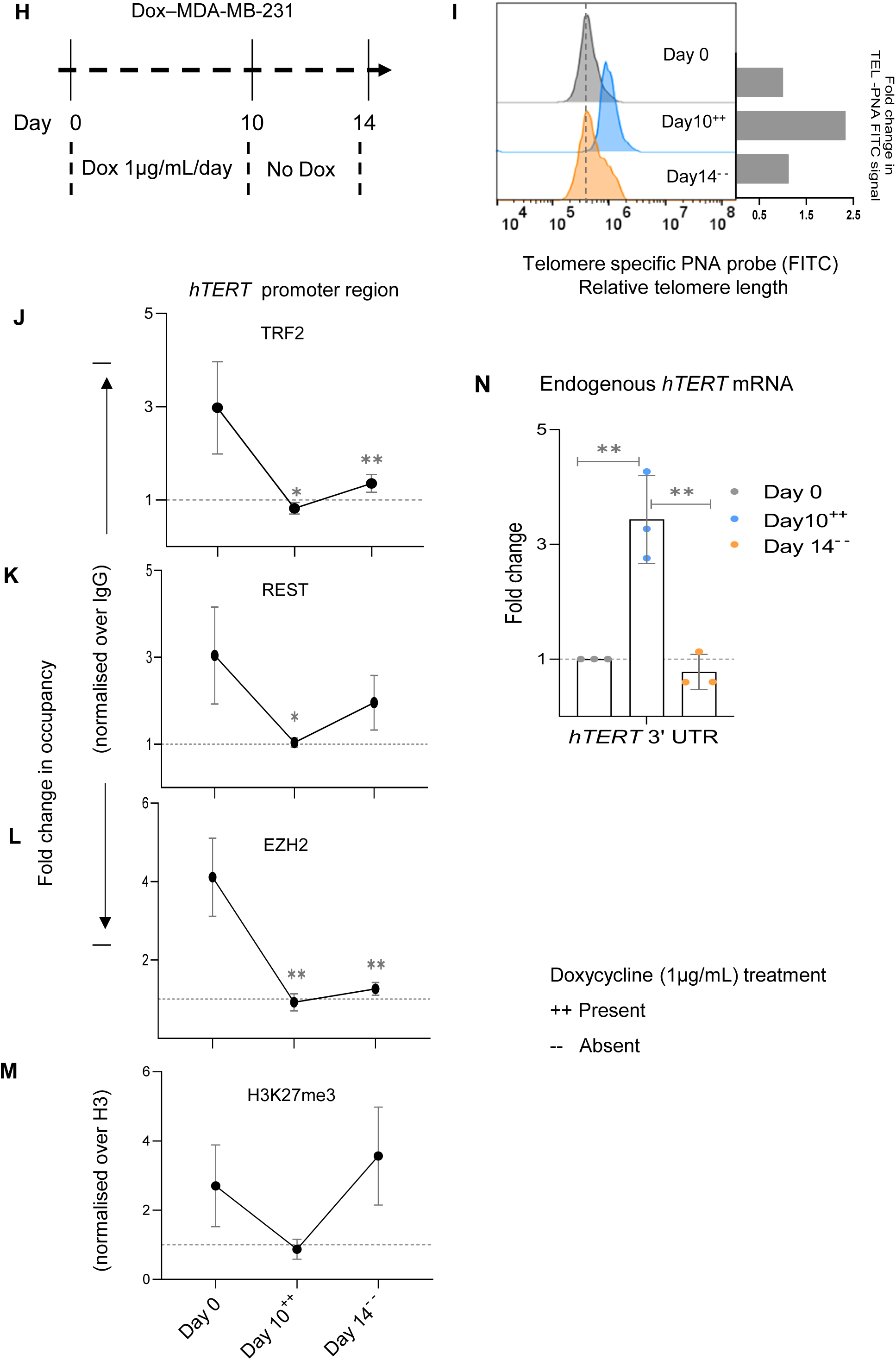
Temporal telomere length elongation followed by shortening shows telomere-sensitive transcriptional regulation of *hTERT* (A) Scheme depicting protocol followed for doxycycline (Dox) inducible hTERT overexpression in HT1080 cells (Dox-HT1080). ++/-- denotes the presence/ absence of dox at the indicated day points. (B) Relative fold change in telomere length at Day 0,10, and 24 determined by FACS in Dox-HT1080 cells; quantification in right panel. (C-F) ChIP followed by qRT-PCR at the 0-300 bp *hTERT* promoter (upstream of TSS) for TRF2 (C), REST (D), EZH2 (E) or H3K27me3 (F) in Dox-HT1080 cells at Day 0,10, and 24; occupancy normalised to respective IgG or total Histone H3 (for H3K27me3). (G) *hTERT* mRNA expression by qRT-PCR using *hTERT* specific 3’UTR primers in Dox-HT1080 cells at day intervals (as indicated); normalised to *GAPDH* mRNA levels. Fold changes were calculated independently for each biological replicate, as the three conditions represent paired samples (uninduced, induced with doxycycline, and post-doxycycline withdrawal). (H) Scheme depicting protocol followed for doxycycline (Dox) inducible hTERT overexpression in MDA-MB-231 (Dox-MDA-MB-231). ++/-- denotes the presence/ absence of dox at the indicated day points. (I) Relative fold change in telomere length at Day 0,10, and 14 determined by Flow cytometry in Dox-MDA-MB-231; quantification in right panel. (J-M) ChIP followed by qRT-PCR at the 0-300 bp *hTERT* promoter (upstream of TSS) for TRF2 (J), REST (K), EZH2 (L) or H3K27me3 (M) in Dox-MDA-MB-231 cells at Day 0,10, and 14; occupancy normalised to respective IgG or total Histone H3 (for H3K27me3). (N) *hTERT* mRNA expression by qRT-PCR using hTERT specific 3’UTR primer in Dox-MDA-MB-231 cells at day intervals (as indicated); normalised to *GAPDH* mRNA levels. Fold changes were calculated independently for each biological replicate, as the three conditions represent paired samples (uninduced, induced with doxycycline, and post-doxycycline withdrawal). Error bars represent ± SDs from the mean of 3 independent biological replicates of each experiment. One-way ANOVA followed by post-hoc tests (Tukey’s HSD) was performed to compare means across time points in Figs C-G, J,-N (*p < 0.05, **p < 0.01, ***p < 0.005, ****p < 0.0001).

TRF2 binding at the *hTERT* promoter decreased at Day 10 with TL elongation. Conversely, as TL shortened beyond Day 10, TRF2 occupancy on the *hTERT* promoter was regained on Day 24; Fig 3C). Notably, this showed TRF2 binding at the *hTERT* promoter to be temporally controlled as a function of TL. Second, this further showed that *hTERT* promoter-bound TRF2 was non-telomeric because telomere-associated TRF2 would increase with TL elongation (and looping), instead of decrease as seen here.

We then examined the occupancy of REST and EZH2 at the *hTERT* promoter at days 0, 10, and 24: decrease in occupancy of both REST and EZH2 on Day 10, relative to Day 0, followed by an increase on Day 24 was clear (Fig 3D and 3E).

Accordingly, H3K27me3 deposition at the *hTERT* promoter was lower on Day 10 compared to Day 0, followed by an increase on Day 24 (Fig 3F). Taken together, clearly REST and EZH2 binding, and H3K27me3 occupancy were as expected from a decrease/increase in TRF2 binding at the *hTERT* promoter as TL increased and then receded.

Consistent with this, expression of the endogenous *hTERT* (using 3’UTR specific qRT-PCR primers) increased by Day 10 from Day 0 levels, and was significantly reduced by Day 24 compared to the Day 10 levels although not reduced to the Day 0 state (Fig 3G), supporting the role of TL elongation in inducing or TL shortening in suppressing *hTERT*.

We considered a second temporal model using stable MDA-MB-231 cells with dox-inducible hTERT to exclude the possibility of cell-type-specific effects. After dox induction, we followed TL elongation/shortening and telomerase activity (through days 4/8/10 and 14, Supplementary Fig 2A-B, right panel; ++/-- denotes presence/absence of dox, respectively). Like HT1080 cells, here also TL elongated till Day 10 and beyond Day 10, the presence of dox showed no further elongation in TL (data not shown). Like the HT1080 cell model dox was withdrawn at Day 10, however, in the case of MDA-MB-231 cells TL shortened to roughly the initial (Day 0) state by Day 14; in a relatively short timeframe than what was noted for HT1080 cells (Supplementary Fig 2A, right panel).

For detailed analysis here we selected days 0, 10 with dox, withdrawal of dox at Day 10 (consistent with the HT1080 cell model), and thereafter Day 14 (given the enhanced reduction in TL in MDA-MB-231 compared to HT1080 cells) as representative time points (Dox-MDA-MB-231); Fig 3H). TL elongated to ∼2-folds by Day 10; and receded to approximately initial (Day 0) length by Day 14 after stopping dox at Day 10 (Fig 3I).

Decrease in TRF2 binding, and corresponding reduced REST, EZH2 and the H3K27me3 mark at the *hTERT* promoter were clear on Day 10 relative to Day 0 (Fig 3J-M). Promoter TRF2, REST and EZH2 occupancy, including H3K27 trimethylation, was regained in the subsequent time point of Day 14, relative to the Day 10 levels, as expected from TL shortening after Day 10 (Fig 3J-M). Accordingly, we noted endogenous *hTERT* transcript followed TL elongation/shortening: increased progressively till Day 10, and subsequently reduced gradually at Day 14 (Fig 3N). Cell-type specific differences in the experimental models, for instance relatively low regain of TRF2 occupancy at the final time day points, Day 14 in MDA-MB-231 compared to Day 24 HT1080 cells, other than relatively enhanced pace of shortening in MDA-MB-231 cells vis-à-vis HT1080 cells on dox withdrawal were interesting to note. Nevertheless, taken together, data from two independent models of temporally altered TL, further supported promoter TRF2-binding mediated *hTERT* regulation to be TL-dependent.

### Telomere-dependent *hTERT* regulation operates *in vivo* within xenograft tumours

We next sought to test TL-sensitive TRF2 binding at the *hTERT* promoter *in vivo.* Xenograft tumours were grown with either HT1080-ST or HT1080-LT cells in NOD/SCID mice (n=5 each, for tumour size characterization see Methods, Fig 4A). Following harvesting, tumours were first characterized: TL was overall higher in all LT tumours compared to HT1080-ST tumours; increased *hTERT* and telomerase activity was retained in HT1080-LT tumours relative to ST (Fig 4B, C; Supplementary Fig 3A). TRF2-ChIP from tumour tissue showed HT1080-ST had significantly more TRF2 occupancy at the *hTERT* promoter than HT1080-LT, consistent with results from cultured cells (Fig 4D and Fig 2B). Correspondingly, H3K27me3 mark deposition was reduced in xenograft HT1080-LT tumours compared to HT1080-ST (Fig 4E) supporting TL-dependent *hTERT* regulation was retained in cells growing *in vivo*.

**Fig 4.**
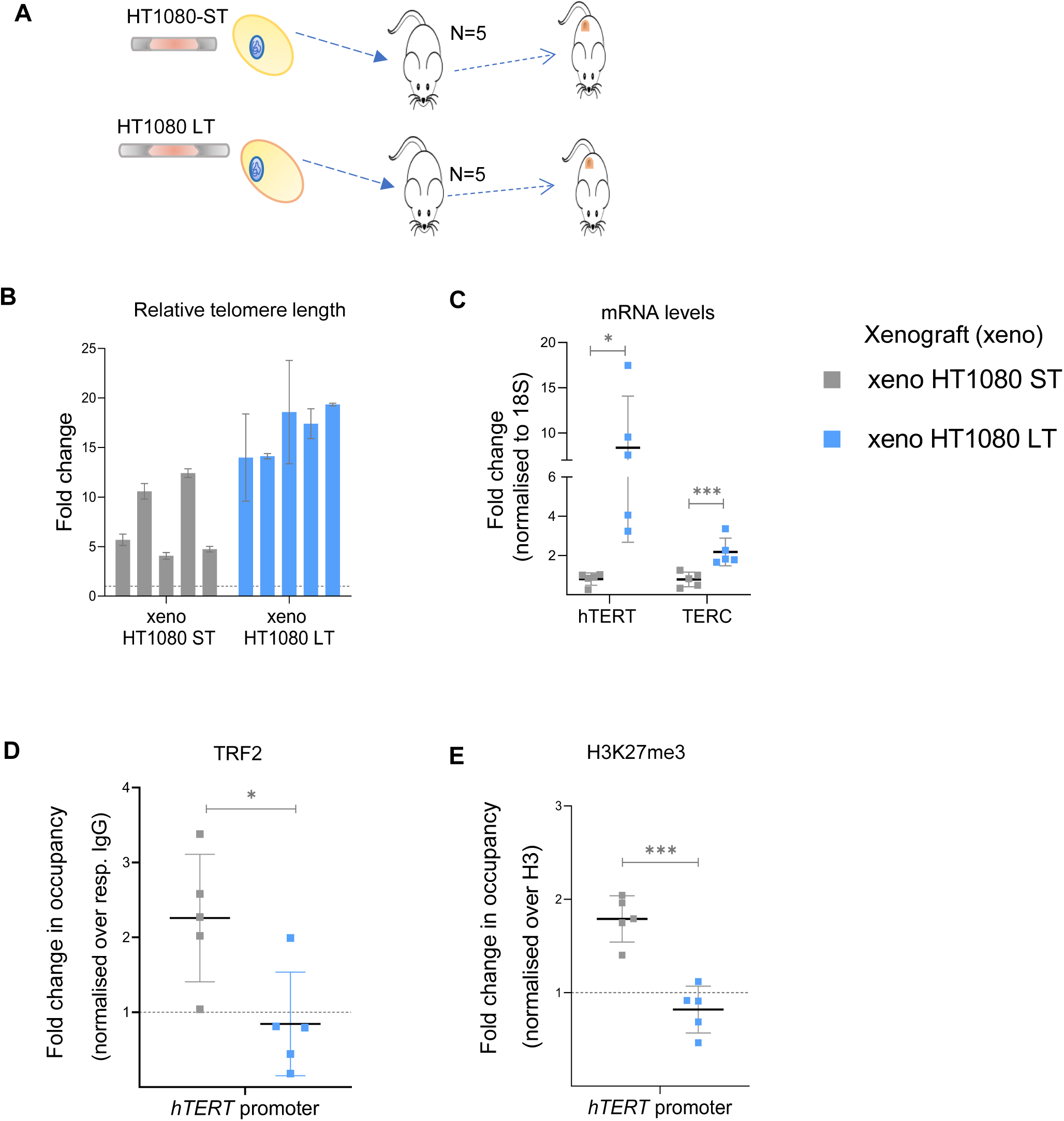
Telomere length sensitive *hTERT* regulation in *in vivo* tumour xenografts (A) Scheme depicting the generation of tumour xenograft with HT1080-ST or HT1080-LT cells. (B) Relative fold change in telomere length in xenograft samples determined by qRT-PCR-based telomere length detection method as reported earlier (O’Callaghan et al., 2008 and Cawthon et al 2002). Telomeric signal normalised over single copy gene, 36B4. (C) *hTERT* (exon 15/16 full-length transcript) and *hTERC* mRNA expression in xenograft tissues by qRT-PCR; normalised to 18S mRNA levels. (D, E) ChIP followed by qPCR at the 0-300 bp *hTERT* promoter (upstream of TSS) for TRF2 (D) and H3K27me3 (E); occupancy normalised to respective IgG and total H3 (for H3K27me3) Error bars represent ± SDs across individual values of n=5 xenograft tumour samples in each group.. p values are calculated by unpaired t-test with Welch’s correction (*p < 0.05, **p < 0.01, ***p < 0.005, ****p < 0.0001).

### G-quadruplex mediated TRF2 binding is essential for telomere-dependent *hTERT* regulation

G-quadruplex (G4) DNA secondary structures(32), widely reported as gene regulatory motifs(33–36), were found in the *hTERT* promoter(37,38). G4-dependent promoter TRF2 binding was also recently shown to regulate *hTERT*(28). Here we asked, whether and how TL-dependent *hTERT* regulation was affected by the promoter G4s. We used the *hTERT* promoter-*Gluc*-reporter inserted at the CCR5 locus in HEK293T cells described above (Fig 2G). G>A mutations at -124 or -146 positions from TSS of *hTERT* promoter are frequently found to be clinically associated with multiple cancers and reported to disrupt the G4s(12,39–44). These mutations were introduced in the reporter individually and a pair of cell lines with either long or short TL were generated (Fig 5A-B).

**Fig 5.**
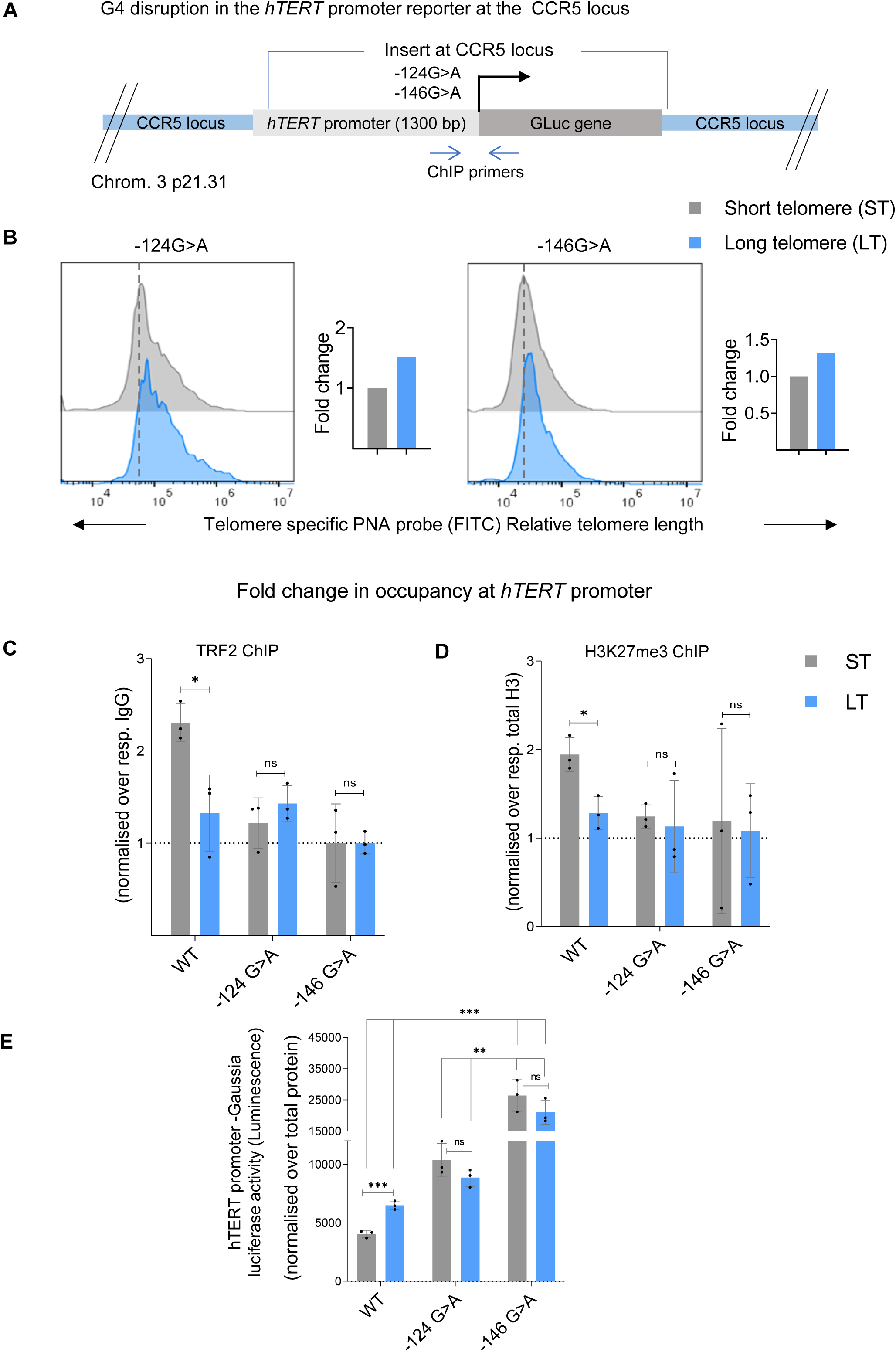
G-quadruplex mediated TRF2 binding is essential for telomere-dependent *hTERT* regulation (A) Scheme depicting CRISPR modified HEK293T cells with 1300 bp *hTERT* promoter with G>A substitution at -124 or -146 bp (upstream of TSS) driving gaussia luciferase (Gluc) construct at the CCR5 safe harbour locus. Scheme denotes ChIP primers used to study chromatin occupancy of 0-300 bp *hTERT* promoter region inserted at the exogenous locus. (B) Relative fold change in telomere length in two independent pairs of cells with short or long telomeres containing *hTERT* promoter G4 disrupting mutations (-124G>A or - 146G>A) at CCR5 locus *hTERT* promoter insert as determined by Flow cytometry; quantification in right panel. (C, D) ChIP followed by qRT-PCR at the inserted *hTERT* promoter (0-300 bp upstream of TSS) for TRF2 (C), and H3K27me3 (D) in pairs of cells generated with -124G>A or -146G>A mutation with long/ short telomeres, along with WT promoter ST/LT pair (as in Figure 2 I,J). Occupancy normalised to respective IgG or total histone H3 (for H3K27me3). (E) *hTERT* promoter-gaussia luciferase activity in short or long telomere cells with - 124G>A and - 146G>A mutated *hTERT* promoter sequence, along with WT promoter ST/LT pair (as in Figure 2 K). Reporter activity presented as luminescence (arbitrary units, a.u.) normalised to respective total protein levels. Error bars represent ± SDs from the mean of 3 independent biological replicates of each experiment. Statistical significance was determined by two-way ANOVA followed by Tukey’s post hoc test for all pairwise comparisons. For planned comparisons between each parental and short-telomere cell line, unpaired t-tests were used. (*p < 0.05, **p < 0.01, ***p < 0.005, ****p < 0.0001).

Low promoter TRF2 binding and reduced H3K27 trimethylation in cells with either -124 or - 146 G>A mutations within the *hTERT* promoter-reporter inserted at the CCR5 locus was clear from earlier work(28). Here, based on low TRF2 binding at the G4-disrupted *hTERT* promoter, we reasoned that TL-dependence of *hTERT* regulation would be affected. TRF2 binding at the *hTERT* promoter with G4-disrupting mutations was not regained upon shortening of TL and was consistent in the case of both -124G>A and -146G>A mutations (Fig 5C, left and right panels). This was in contrast to the increase in promoter TRF2 binding observed on TL shortening in the case of the unmutated promoter (Fig 2I), supporting the role of G4s in *hTERT* promoter TRF2 binding and thereby in TL-dependent *hTERT* regulation. As expected, the difference in H3K27me3 deposition and *Gluc* activity was insignificant in -124G>A or -146G>A ST cells compared to the corresponding LT cells (Fig 5D, E).

### TRF2 R17 residue is required for the repression of *hTERT* expression

To understand the role of TRF2 post-translational modification(s) (PTM), if any, on *hTERT* repression we screened TRF2 mutants(45). TRF2 mutants (arginine methylation deficient R17H(46); acetylation deficient K176R, K190R(47) or phosphorylation deficient T188N(48) were expressed in HT1080 cells in a background where endogenous TRF2 was silenced (Supplementary Fig 4A). As expected, TRF2 silencing induced *hTERT*, and re-expression of TRF2-wild type (WT) repressed *hTERT*. Interestingly, R17H and K176R increased *hTERT*, whereas T188N and K190R gave relatively moderate change (Supplementary Fig 4B). Noting that the N-terminal domain of TRF2 (with the R17 residue) was shown to interact with G4 DNA(32,49), and also the core histone H3(50) we focused on understanding the role of TRF2-R17 at the *hTERT* promoter.

Dox inducible TRF2-R17H or TRF2-WT stable lines were made with HT1080, HCT116 and MDA-MB-231 cells. Induction of TRF2-R17H gave significant upregulation of *hTERT*; whereas as expected TRF2-WT repressed *hTERT* in all three cell lines (Fig 6A, Supplementary Fig 4C; dox dose-dependent TRF2 induction and resultant *hTERT* levels shown in Supplementary Fig 4D). The increase in *hTERT* with TRF2-R17H expression in contrast to its reduction with TRF2-WT was also clear from *hTERT* FISH (Fig 6B).

**Fig 6.**
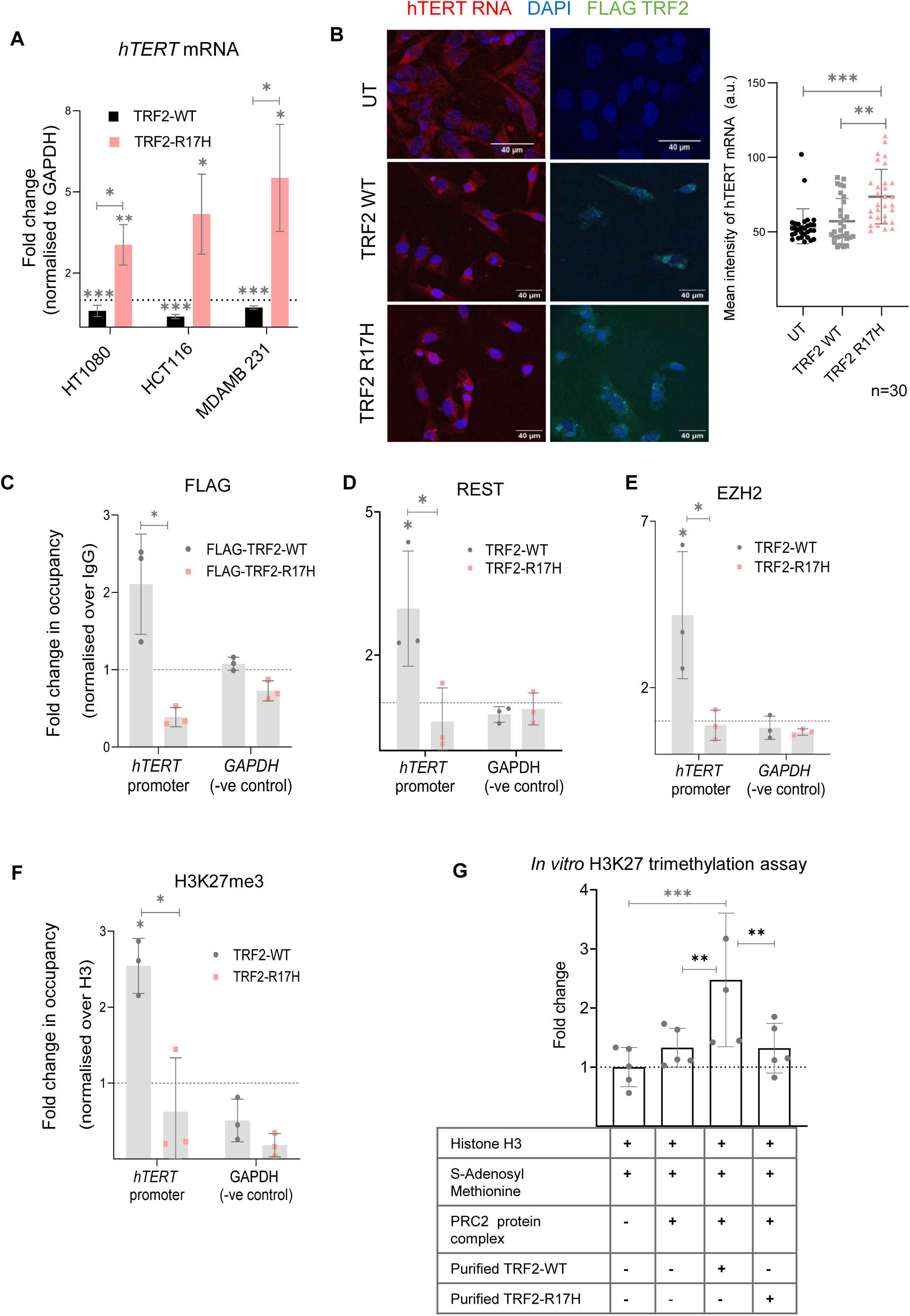
TRF2 R17 residue is required for the repression of *hTERT* expression (A) *hTERT* full-length transcript (exon 15/16) mRNA expression levels by qRT-PCR; normalised to *GAPDH* mRNA levels upon stable doxycycline induction of wildtype (WT) TRF2 or R17H TRF2 mutants in HT1080, HCT116 or MDA-MB-231 cells (B) *hTERT* mRNA FISH upon TRF2 WT or TRF2 R17H overexpression (un-transfected, UT as control) in HT1080 cells; quantification shown in right panel. FLAG-tagged TRF2 overexpression was confirmed by Immuno-fluorescence microscopy. (C-F) ChIP followed by qRT-PCR at the 0-300 bp *hTERT* promoter (upstream of TSS) for FLAG-tagged TRF2 (C), REST (D), EZH2 (E) or H3K27me3 (F) in HT1080 cells upon expression of WT TRF2 or TRF2 R17H. Occupancy normalised to respective IgG and total Histone H3 (for H3K27me3); qRT-PCR on the *GAPDH* promoter was used as the negative control in all cases. (G) In vitro methyl transferase activity of the reconstituted PRC2 complex resulting in H3K27 trimethylation in the presence or absence of TRF2 WT or TRF2 R17H protein. Error bars represent ± SDs from the mean of 3 independent biological replicates of each experiment. Unpaired t-tests were conducted to assess the significance of each condition individually within the same cell line, and to compare the two conditions TRF2-WT or TRF2-R17H in (A); one-way ANOVA followed by post-hoc tests (Tukey’s HSD) was performed to compare means across the three conditions in (B); p values are calculated by unpaired t-test in (C-F); and two-way ANOVA followed by post-hoc tests (Tukey’s HSD) in (G). (*p < 0.05, **p < 0.01, ***p < 0.005, ****p < 0.0001).

To understand loss of function we looked closely at TRF2-R17H DNA binding at the *hTERT* promoter. The binding of FLAG tagged-TRF2-R17H on the *hTERT* promoter was insignificant compared to the FLAG tagged-TRF2-WT in HT1080 cells (Fig 6C). As expected from the loss of TRF2-R17H binding, REST, EZH2 and H3K27me3 occupancy was significantly low on the *hTERT* promoter for TRF2-R17H compared to TRF2-WT (Fig 6D-F).

### TRF2 promotes H3K27 trimethylation *in vitro*

For a deeper understanding of the role of TRF2 in H3K27 trimethylation, we used *in vitro* histone H3 methyltransferase assay. Purified histone H3 along with the reconstituted PRC2 repressor complex was analyzed in the presence/absence of recombinant TRF2-WT or TRF2-R17H for methyltransferase activity (Fig 6G, Supplementary Fig 4E-F). H3K27 trimethylation in the presence of H3 and the PRC2 complex was first confirmed. Following this, treatment with purified TRF2-WT gave further increase in H3K27 trimethylation whereas TRF2-R17H did not lead to any significant increase, relative to respective controls (Fig 6G). Interestingly, together these support a direct function of TRF2 in histone H3K27 trimethylation in presence of the PRC2 complex, where the R17 residue of TRF2 plays a necessary role.

### Telomere length regulates *hTERT* expression in induced pluripotent stem cells (iPSCs)

To test the effect of TL on *hTERT* expression in a physiological setting where telomeres elongate or shorten, we used fibroblast cells along with corresponding induced pluripotent stem cells (iPSCs). Primary foreskin (FS) fibroblasts were reprogrammed to pluripotent stem cells (iPSCs) (see Methods, scheme in Fig 7A, characterization data for iPSC in Fig 7B). *hTERT* mRNA expression (Fig 7C), telomerase activity (Fig 7D) and relative telomere length (Fig 7E), were confirmed, and as expected were higher in the iPSCs compared to parent FS fibroblast cells. Chromatin binding of TRF2, REST, and EZH2 proteins and deposition of H3K27me3 mark were analysed in FS fibroblast and derived iPSCs (Fig 7F-I). TRF2 occupancy in the iPSCs with longer telomeres was found to be lower on the *hTERT* promoter in comparison to parent fibroblasts (Fig 7F). Further, occupancy of REST (Fig 7G) and the repressor complex protein, EZH2 (Fig 7H) was reduced in the pluripotent stem cells. Corresponding to the reduced levels of TRF2 occupancy and resultant lower binding of the REST/PRC2 complex on the *hTERT* promoter in the iPSCs, the repressor mark, H3K27me3 was significantly lower relative to FS Fibroblast cells (Fig 7I).

**Fig 7.**
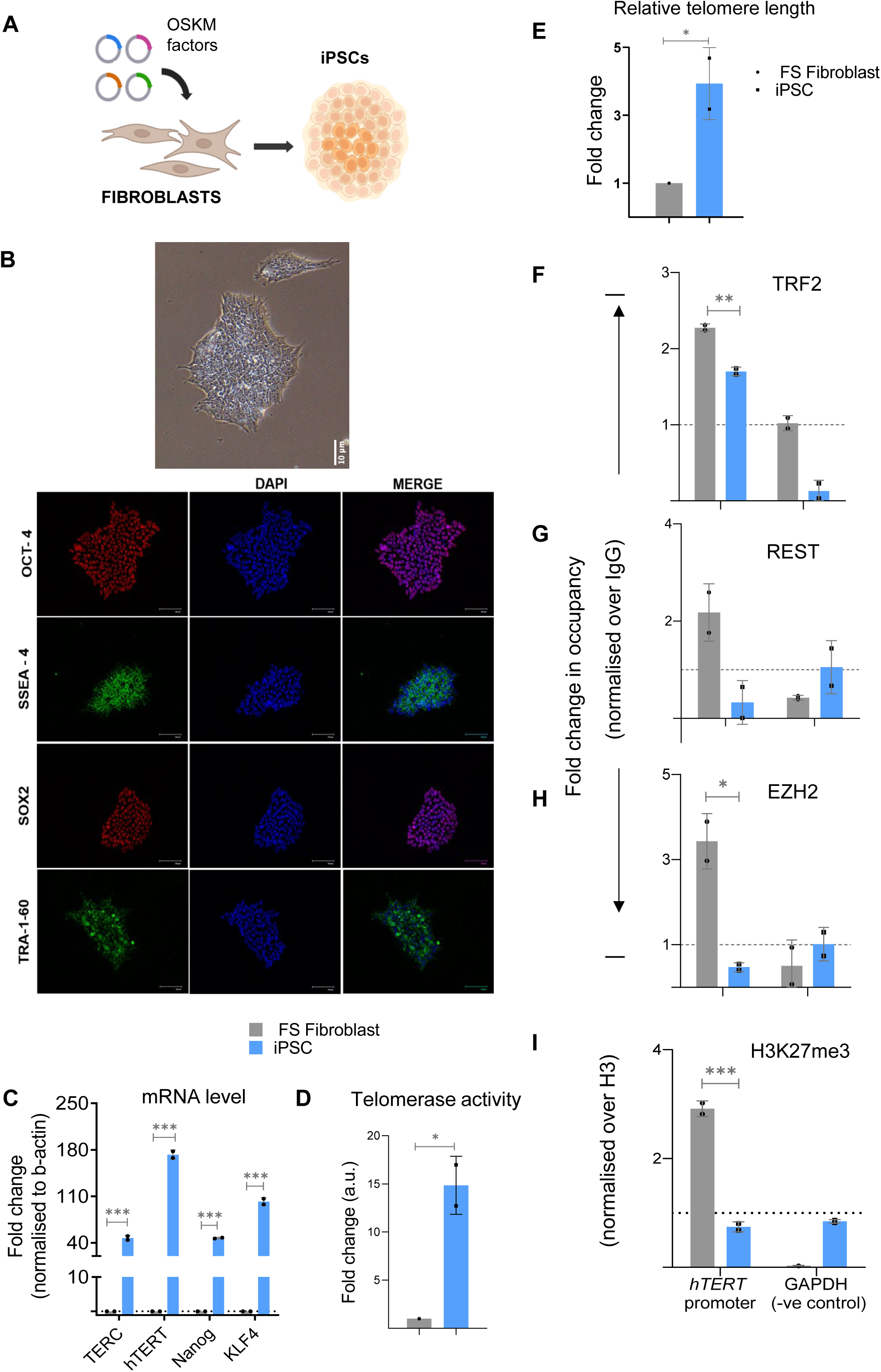
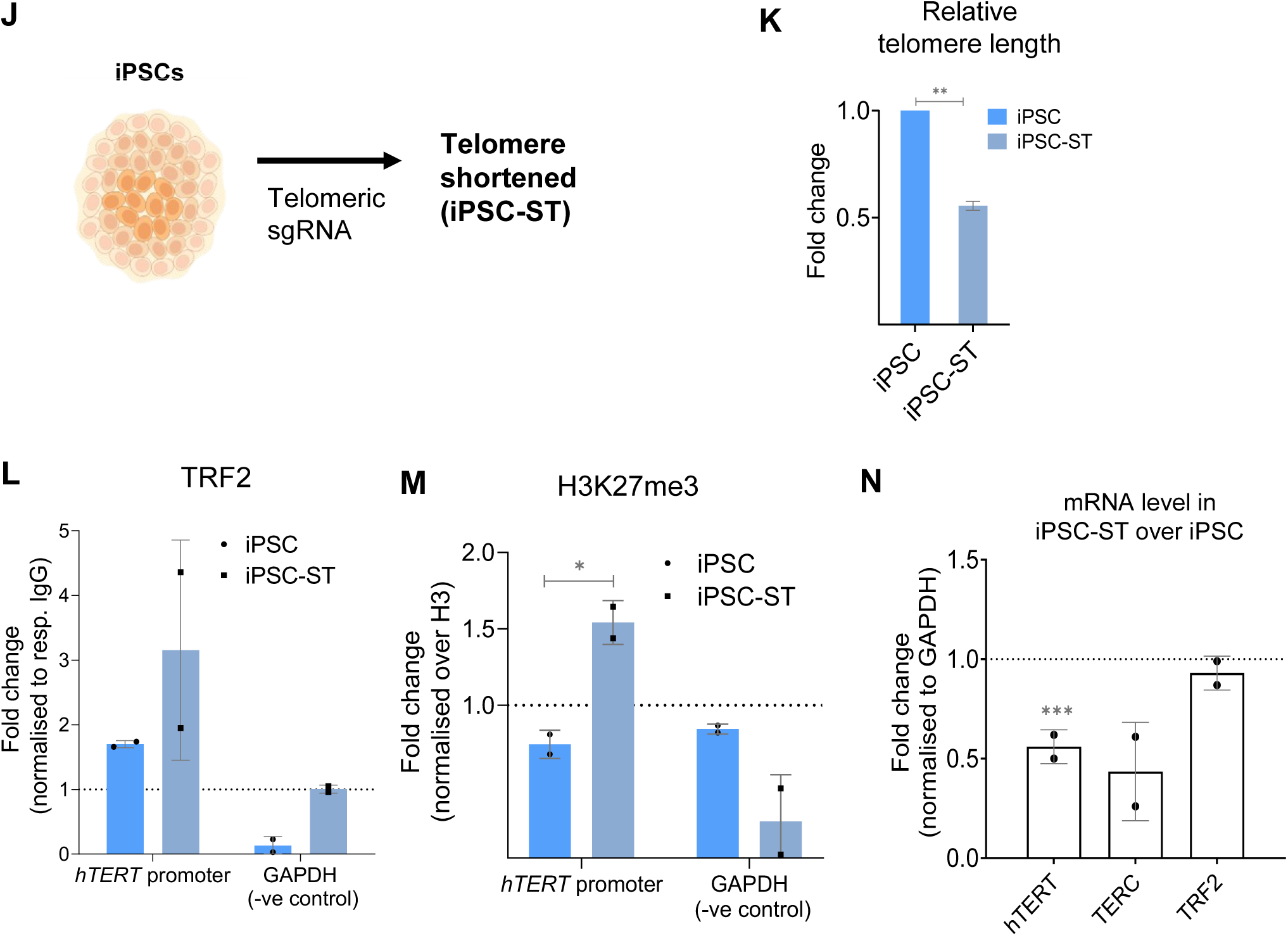
Telomere length regulates *hTERT* expression in iPSC (A) Scheme showing generation of induced pluripotent stem cells (iPSCs) from foreskin fibroblast (FS Fibroblast) cells by overexpressing Yamanaka factors (Oct4, Sox2, Klf4, Myc) (B) Characterization of iPSCs (Upper panel, bright field image) generated from FS Fibroblast cells by immuno-fluorescence using Oct-4, SSEA-4, Sox-2 and TRA-1-60 antibodies as stemness markers. (C) mRNA levels for *hTERT* (full-length exon 15/16 transcript), TERC (RNA component) and stemness marker genes Nanog, Klf4 in FS fibroblast and derived iPSC, analysed in pairs in each biological replicates. (D) Telomerase activity in FS Fibroblast cells and derived iPSCs determined using telomerase-repeat-amplification-protocol (TRAP) followed by ELISA (see Methods); (E) Relative fold change in telomere length in primary FS Fibroblast cells and derived iPSC, determined by qPCR-based telomere length detection method (F-I) ChIP followed by qPCR at the 0-300 bp *hTERT* promoter (upstream of TSS) for TRF2 (F), REST (G), EZH2 (H) and H3K27me3 (I) up to 300 bp upstream of transcription start site (TSS); occupancy normalized to respective IgG or total Histone H3 (for H3K27me3). qPCR on the GAPDH promoter was used as a negative control in all cases. (J) Scheme depicting generation of induced pluripotent stem cells (iPSCs) with shortened telomeres, (iPSC-ST) using telomere-specific sgRNA guided CRISPR-Cas9 to trim telomeres. (K) Relative fold change in telomere length in iPSC-ST cells with respect to unaltered iPSC, determined by qPCR-based telomere length detection method (L-M) ChIP followed by qPCR at the 0-300 bp *hTERT* promoter (upstream of TSS) in iPSC-ST ells in comparison to unaltered iPSC, for TRF2 (L) and H3K27me3 (M); occupancy normalized to respective IgG or total Histone H3 (for H3K27me3). qPCR on the GAPDH promoter was used as a negative control in all cases. (N) mRNA levels for *hTERT* (full-length exon 15/16 transcript), TERC (RNA component) and TRF2 in iPSC-ST over unaltered iPSC. All error bars represent ± SDs from the mean of 2 independent biological replicates of each experiment. p values are calculated by unpaired t-test (*p < 0.05, **p < 0.01, ***p < 0.005, ****p < 0.0001).

Next, we asked if TL played a causal role in *hTERT* expression. That is, whether TL change in iPSCs influenced TRF2 occupancy at the *hTERT* promoter and its subsequent impact on *hTERT*. We used the telomere-specific sgRNA-guided CRISPR-Cas9 to trim telomeres (as reported earlier) (51). This resulted in a population of iPSCs with short telomeres (’iPSC-ST’; ∼50% reduction in TL; see Methods, Fig. 7J, K). A comparison of TRF2 occupancy between iPSC-ST cells and unaltered iPSCs revealed a marked increase in TRF2 binding at the *hTERT* promoter (∼3-fold) in the telomere-shortened cells (Fig. 7L). As expected from this based on the above findings, we observed an increase (of 1.5-fold) in the deposition of the repressive histone mark H3K27me3 in iPSC-ST cells compared to unaltered iPSCs (Fig. 7M). Further, consistent with these, *hTERT* was significantly reduced in iPSC-ST cells relative to unaltered iPSCs (Fig. 7N). Taken together, these demonstrate how *hTERT* repression results of TL shortening through non-telomeric TRF2.

## Discussion

Relatively long telomeres induced permissive chromatin at the *hTERT* promoter. Conversely, shorter telomeres led to closed chromatin at the promoter (Fig 2E-F, Fig 3D, E, K, L). This resulted in telomere-dependent *hTERT* transcription in multiple cell types/models as in: (a) short/long telomere HT1080, HCT116, MDA-MB-231 cells, and primary FS fibroblast along with corresponding derived iPSC; (b) temporal TL elongation with subsequent shortening; (c) TL-dependent transcription from artificially inserted *hTERT* promoter; (d) and, tumour cells grown *in vivo* in mice. Mechanistically, TRF2 binding at the *hTERT* promoter decreased/increased with TL elongation/shortening respectively, affecting TRF2-dependent recruitment of epigenetic modulators REST and the PRC2 repressor complex (Fig 2-7). The resulting loss/gain in the repressor histone H3K27 trimethylation up or down-regulated *hTERT* respectively, as is shown in the following scheme (Fig 8). This was consistent with the TSP model described by us earlier (Fig 1) (13,21,27).

**Fig 8.**
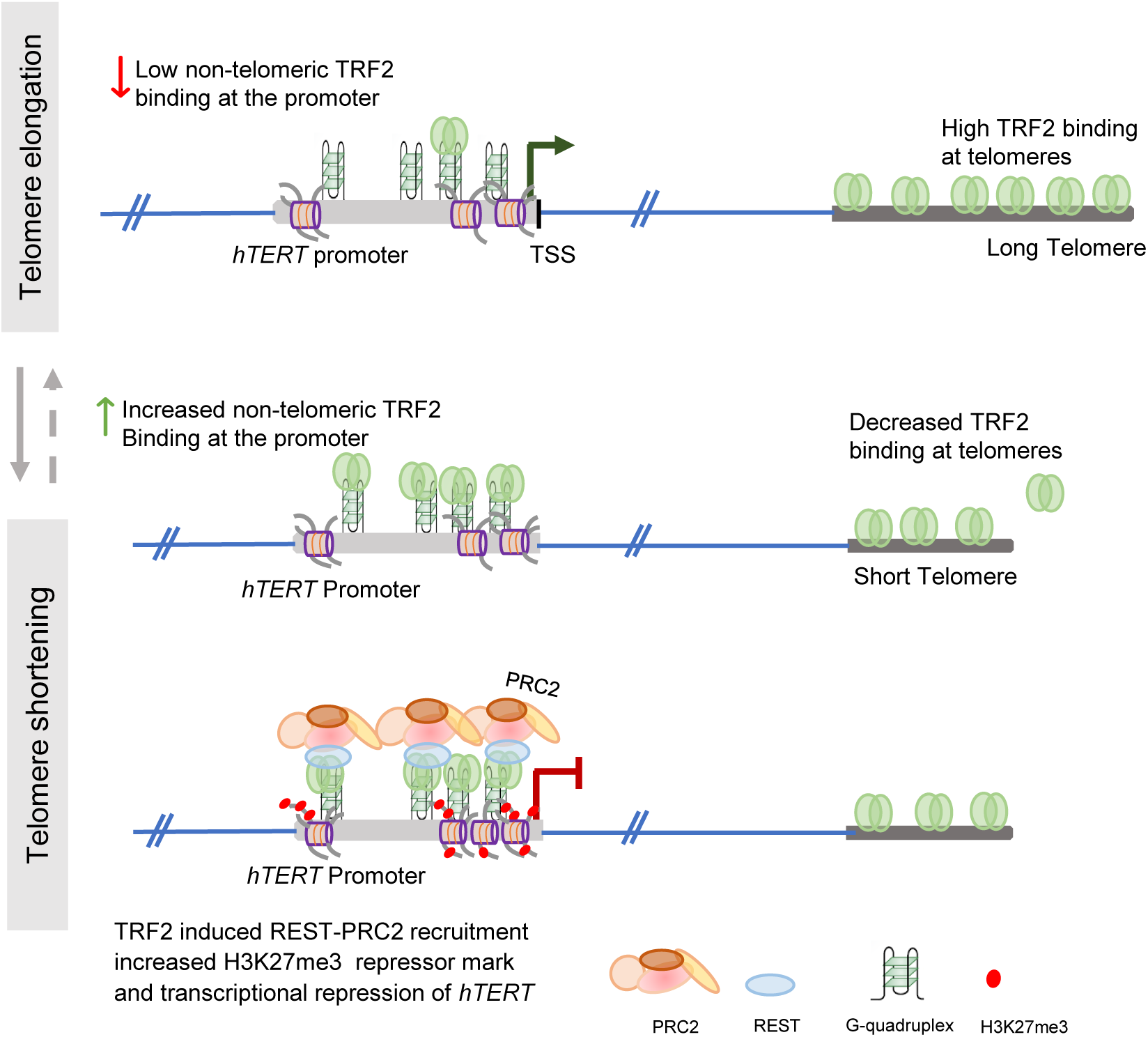
Telomere length controls human Telomerase (*hTERT*) expression through non-telomeric TRF2 Illustration of the telomere-dependent epigenetic modification of chromatin structure at the *hTERT* promoter, resulting in upregulation or downregulation of *hTERT* expression due to altered non-telomeric TRF2 binding at the promoter. Relatively long telomeres cells (top panel) have lower TRF2 binding at the *hTERT* promoter, promoting permissive chromatin and upregulation of *hTERT* transcription. Conversely, shorter telomeres cells (bottom panel) with increased TRF2 binding at the *hTERT* promoter recruit more REST-PRC2 epigenetic complex causing increased repressor histone H3K27 trimethylation deposition. This leads to a more closed chromatin state at the *hTERT* promoter, suppressing its transcription.

The overwhelming understanding, particularly in cancer cells, suggests that the reactivation of *hTERT* drives telomere elongation/maintenance(3,9,14,52). On the other hand, however, the possibility that telomeres conversely might impact the regulation of *hTERT* remains unclear. To test this, here we focused on two contexts: cancer cell lines engineered to modify telomere length (TL) and cellular reprogramming where TL changes are evident (53,54). Multiple lines of evidence support the causal role of telomeres in *hTERT* regulation. (a) TL elongation, by exogenous *hTERT*, or independent of *hTERT* (using G-rich telomeric oligonucleotides as reported earlier (see Methods)) in different cell types produced *hTERT* upregulation (Fig 2). (b) TL shortening, on the other hand, caused *hTERT* suppression (Fig 2). (c) Temporally induced TL elongation over days, using exogenous *hTERT*, gradually activated the endogenous *hTERT* promoter. As TL receded, after discontinuing induction, the endogenous *hTERT* promoter was re-suppressed (Fig 3). (d) *hTERT* activation during reprogramming of primary fibroblast to iPSCs showed reduced promoter TRF2 binding on telomere elongation and *hTERT* activation (Fig 7). (e) And, finally upon telomere shortening in iPSCs, TRF2 occupancy was regained on the *hTERT* promoter, followed by increased repressor mark deposition and reduced *hTERT* transcription. Together, these demonstrate that telomeres play a key role in *hTERT* regulation, adding new insight to the current understanding of telomere elongation/maintenance as a consequence of hTERT activity (9,55,56).

Telomere length and *hTERT* expression are critical in maintaining pluripotency and the ability of stem cells to differentiate (53,54,57,58). The re-occupation of TRF2 at the *hTERT* promoter in telomere-shortened iPSCs (iPSC-ST) along with suppression of its expression demonstrates the applicability of TRF2-dependent TL-sensitive *hTERT* regulation in stem cells and presents a potentially useful tool to modulate its differentiation capacity and pluripotency with further exploration.

An earlier paper showed long telomeres on chromosome 5p interacted with the proximal *hTERT* locus (1.2 Mb away) by physical chromatin looping (TPE-OLD)(19). Resulting heterochromatinization due to telomeric factors, including TRF2, repressed *hTERT*; loss of looping on telomere shortening de-repressed *hTERT*. We, on the other hand, observed that long telomeres induce, and shorter telomeres repress *hTERT* transcription, suggesting this was distinct from looping-induced interactions. An important underlying difference in the nature of TRF2 at the *hTERT* promoter must be noted: in TPE-OLD, TRF2 is telomere-associated, whereas TRF2 is non-telomeric in TSP. To confirm non-telomeric TRF2 binding we reasoned in two ways. (a) Telomere-associated TRF2 is likely to be present with other telomeric factors like RAP1 and TRF1(21), and (b) inserted an *hTERT* promoter-reporter >40 Mb from telomeres where looping interaction was unlikely. While TRF2 binding at the *hTERT* promoter was clear, RAP1 or TRF1 was absent(28). Furthermore, *hTERT* promoter activity from the inserted reporter was TL-dependent (Fig 2G-K). These support *hTERT* promoter-bound TRF2 as non-telomeric and indicate the two models (TPE-OLD and TSP) are likely context-specific. However, additional experimentation would be required to ascertain the contexts or range of telomere length at which the two mechanisms could operate in isolation or concert – providing further insights into their physiological relevance.

*hTERT* promoter G4s had a causal role in telomere-induced *hTERT* regulation (Fig 5). Interestingly, in many cancers, activation of *hTERT* was reported to be frequently due to *hTERT* promoter mutations; subsequently noted to overlap with the *hTERT* promoter G4 forming stretch (12,13,37–41,43,44). We show for the two most common clinically significant *hTERT* promoter mutations known to abrogate these G4 structures, the TRF2 binding site on the *hTERT* promoter was also disrupted (Fig 5C) (28). The artificially introduced *hTERT* promoter-reporter on G4 disruption lost its dependence on TL (Figs 2 and 5). These build support for a G4-dependent TL-driven mechanism of *hTERT* regulation/maintenance (11,59).

Xenograft mice tumours indicated TL-induced *hTERT* activation might be active *in vivo*. However, results are primarily from cells engineered for TL elongation/shortening (Fig 4). While these open new avenues additional work would be necessary to understand broader implications. For example, it is not clear how the initial TL elongation (which results in *hTERT* reactivation) is triggered during tumorigenic transformation. Interestingly, recent work shows telomere transfer from antigen-presenting cells (APCs) results in telomere elongation of specific T cell populations, subsequent telomerase expression induced proliferative expansion during T cell activation(60). This suggests the possibility of telomere transfer between tumour and immune/other cells within the tumour microenvironment might alter TL, contributing to *hTERT* activation. A similar model might be functional during reprogramming. Moreover, it would be of interest to test how TL-dependent *hTERT* regulatory mechanisms affect senescing primary cell and telomerase-negative cancer cells (with ALT mechanism).

The current study provides experimental evidence that TRF2, a well-characterized telomere-binding protein, mediates crosstalk between telomeres and the regulatory region of the hTERT gene in a telomere length-dependent manner. Given the observed link between hTERT expression and telomere length, it is likely that additional telomere-associated proteins and regulatory pathways contribute to this regulation.

The remaining shelterin complex components—POT1, hRap1, TRF1, TIN2, and TPP1— may play crucial roles in this context, as they are integral to telomere maintenance and protection(61). Additionally, several DNA damage response (DDR) proteins, which interact with telomere-binding factors and help preserve telomere integrity, could potentially influence hTERT regulation in a telomere length-dependent manner(62). However, direct interactions or regulatory roles would require further experimental validation. Another group of proteins with potential relevance in this mechanism are the sirtuins, which directly associate with telomeres and are known to positively regulate telomere length, undergoing repression upon telomere shortening(63,64). Notably, SIRT1 has been reported to interact with telomerase (65), while SIRT6 has been implicated in TRF2 degradation (47) and telomerase activation(66). Given their roles in telomere homeostasis, sirtuins may serve as key mediators of telomere length-dependent hTERT regulation.

In summary, results reveal a heretofore unknown mechanism of *hTERT* regulation by TRF2 in a TL-dependent manner. Long telomeres sequester more TRF2 for telomere protection, which reduces TRF2 at the *hTERT* promoter. Resulting in de-repression of *hTERT*, and increased *hTERT* levels in turn synthesize (Fig 2) and help maintain longer telomeres. The reversal of this was observed in TL shortening. Taken together, these suggest a feed-forward model mechanistically connecting telomeres to *hTERT* with likely significant implications in better understanding telomere-related physiological events, particularly tumorigenesis, ageing and de-differentiation (14,27,55,67–69).

## Materials and Methods

### Cancer cell lines

HT1080 (obtained from ATCC) and its derivative cell line HT1080-LT were maintained in Minimum Eagle’s Essential Medium (MEM) (Sigma-Aldrich) with 10% Fetal Bovine Serum (FBS) (Gibco). HCT116 p53 null (gift from Prof. Bert Vogelstein’s lab), HCT116 WT, MDA-MB-231 and HEK293T (purchased from NCCS, Pune), CRISPR CCR5 locus insert cell lines; along with their derivative cells (short or long telomeres) were grown in Dulbecco’s Modified Essential Medium High Glucose (DMEM-high Glucose) (Sigma-Aldrich) with 10% FBS. Doxycycline (dox) Inducible TRF2 cells in HT1080 and Dox-HT1080 were cultured in MEM and dox Inducible TRF2 HCT116 and MDA-MB-231 and Dox-MDA-MB-231 cells were cultured in DMEM-High Glucose, all supplemented with 10% tetracycline-free Fetal Bovine Serum (FBS) (Clontech and DSS Takara). Cells were trypsinized and subcultured in desired culture vessels when at 90% confluency. All cells were grown in 5% CO2, 95% relative humidity and 37°C culturing conditions.

### Generation of short or long-TL cellular model systems

1. A longer telomere version of HT1080 fibrosarcoma cells (termed super-telomerase and referred to as HT1080-LT in the current paper) was generated by overexpression of telomerase (hTERT) and telomerase RNA component (*hTERC*) in the HT1080 unmodified cell line (termed as HT1080-ST in the current paper). It was received as a generous gift from Lingner Lab (29). The two cell lines (HT1080 ST/LT) have been cultured and maintained independently of each other.
2. HCT116 p53 null or knockout cells (termed LT here) were used as the parent cell line to generate another stable cell line with short telomere using commercial *hTERC* knockdown plasmid (ST) and a scrambled control (LT) from Santa Cruz Biotechnology. Over multiple passaging under puromycin selection, telomere length reduction was achieved. This HCT116 ST/LT cell line pair was generated under p53 knockout (p53^-/-^) background to prevent p53-regulated *Siah1* mediated TRF2 degradation as reported earlier (70,71).
3. A long telomere length version of the MDA-MB-231 cell line was generated using the alternative lengthening of telomeres (ALT) mechanism(30). MDA-MB-231 cells were seeded and treated with GTR (guanine-rich terminal repeats) oligos at 3uM concentration, in serum-free media for 24h. Post-treatment, the cells were kept in DMEM-High Glucose for 24 hrs. Sequential treatment for 6 feedings (i.e., 12 days), increased telomere length by 4-fold, as determined by Telomeric PNA-Flow cytometry (FACS), compared to its corresponding unmodified parental cell (MDA-MB-231 ST), being cultured simultaneously. These MDA-MB-231 ST/LT cells have been analysed as paired sets in all experiments
4. HCT116 cells were utilised to generate a short telomere length version utilizing CRISPR-Cas9-based telomere trimming. A telomeric sequence-specific sgRNA (tel-sgRNA) and *Sp*Cas9 expressing plasmid were transfected into the cells to generate a telomere-shortened version (ST) of the cells. This mode of telomere shortening is quicker than that via *hTERC* knockdown (as in point 2). Telomere length was quantified by FACS in comparison to the untransfected control HCT116 cells, after 3 days fromr transfection - post puromycin treatment (1µg/mL) to select out transfected cells (with shorter telomeres).These HCT116 ST/LT cells (by telomere trimming) have been analysed as paired sets in all experiments. Telomeric sgRNA: GTTAGGGTTAGGGTTAGGGTTA(51)
5. To generate the TL elongation followed by shortening model, the doxycycline-inducible hTERT expression system, (stable cell generation discussed below), was used. The doxycycline-inducible system consists of cells with an integrated dox-hTERT cassette, seeded in different culture flasks and maintained under three conditions: uninduced (no doxycycline), induced with doxycycline for 10 days, and post-induction withdrawal for defined time periods. Cells are harvested at multiple time points and analysed as paired samples. Briefly, the Dox-HT1080 cells were seeded in T25 flasks and treated with 1µg/mL dox daily (one set being cultured without induction) and TL measured at time points Days 0/6/8/10/16 and 24. Beyond the first 10 days, dox treatment produced no further elongation (as seen on Day 16^--^ in Supplementary Fig 2A left panel). Cells were cultured till Day 24, after discontinuing dox at Day-10, for telomere shortening. Similar treatment was done for the Dox-MDA-MB-231 cellular model, and TL was measured on Days 0/4/8/10/12 and 14; cells were studied till Day 14 (after dox withdrawal at Day-10 and no further increase in TL with dox treatment) when substantial telomere shortening was noted.
6. 1300 bp region of *hTERT* promoter starting from 48 bp downstream of TSS with Gaussia Luciferase reporter was procured from Genecopoeia-HPRM25711-PG04 (pEZX-PG04.1 vector) and inserted in HEK293T cells via CRISPR-Cas9 technology as discussed in(28). The telomeric sequence specific sgRNA (tel-sgRNA) and *Sp*Cas9 expressing plasmid as mentioned for HCT116, these CRISPR modified cells were transfected to generate a telomere shortened version of the CRISPR cells for both unmutated *hTERT* promoter and G-quadruplex (G4) disrupting mutant promoter forms, namely -124G>A and -146G>A. Telomere length was quantified by FACS, after 3 days of transfection post puromycin treatment to select out transfected cells (with shorter telomeres).
7. Reprogramming was conducted using the CytoTune iPS 2.0 Sendai Reprogramming Kit (Thermo Scientific) according to the manufacturer’s instructions. In brief, fibroblast cells (5 × 10^5) were transduced with reprogramming factor genes delivered by non-integrating Sendai viruses. The following day, cells were centrifuged at 200g for 5 minutes at room temperature to remove the virus and then cultured for an additional two days. On the third day post-transduction, 0.5–2 × 10^5 cells were plated on Matrigel (Corning)-coated 6-well plates and maintained in Essential 8™ Medium (Gibco™). The medium was changed daily, and cells were monitored for the emergence of colonies resembling embryonic stem (ES) cells. Between days 16 and 20 following transduction, colonies with flat, well-defined margins indicative of an ES-like phenotype were manually selected and expanded as induced pluripotent stem cells (iPSCs). These iPSC colonies were cultured on Matrigel-coated plates in Essential 8™ Medium at 37°C in a 5% CO2 environment, with daily media changes. During passaging, the typical split ratio was 1:5, and colonies were detached using ReLeSR (Stem Cell Technologies). To generate the telomere shortened version of iPSCs, iPSCs were seeded in Matrigel-coated 6-well plates and transfected with a telomeric sequence-specific sgRNA (tel-sgRNA) and *Sp*Cas9 expressing plasmid. 24h post-transfection, they were subjected to puromycin selection (0.4µg/mL) for 5 days and monitored for any morphological or proliferation capacity changes.

Reagents catalog no:

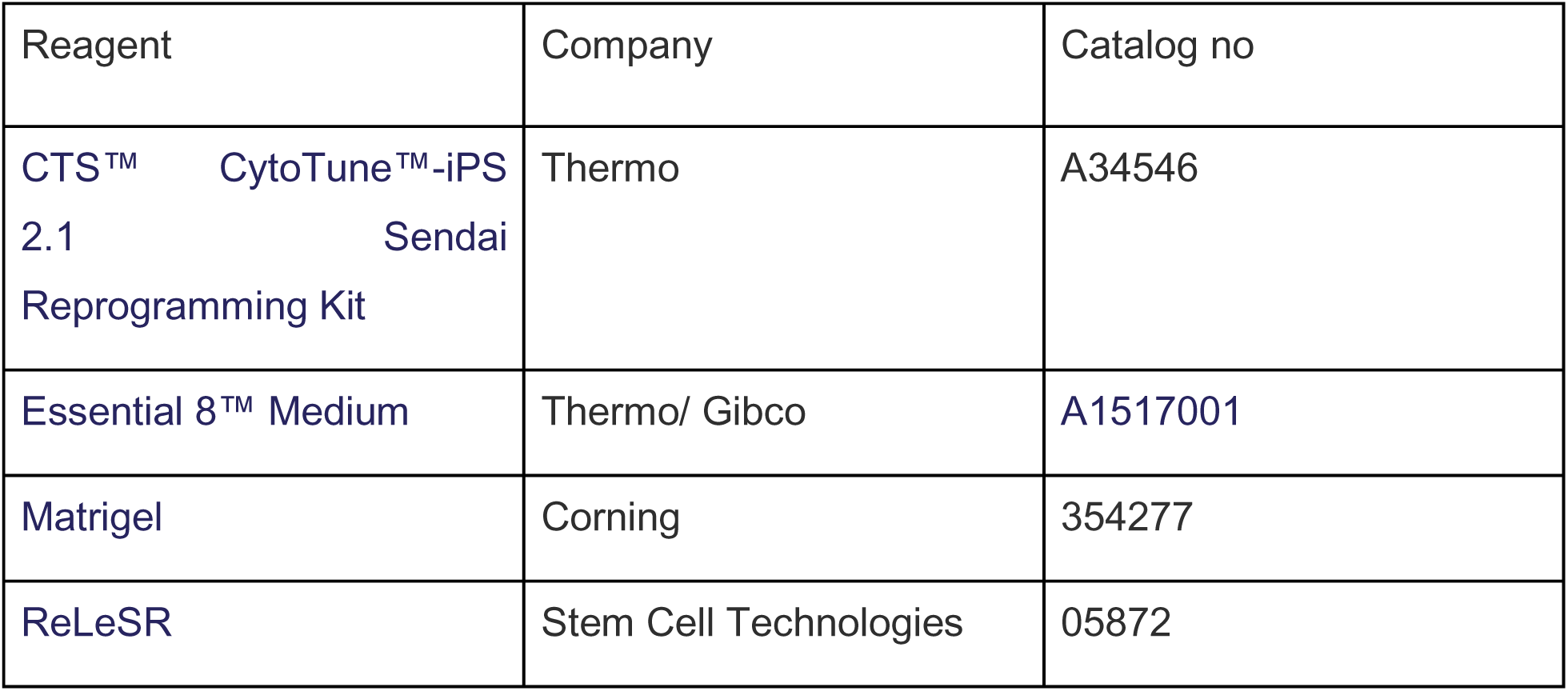

### Doxycycline (dox) inducible hTERT and TRF2 post-translationally-modified (PTM) lentiviral stable cell generation

Doxycycline (dox) inducible hTERT construct and TRF2 WT and R17H PTM variant construct-both in pCW57.1 backbone were individually transfected with 3^rd^ generation lentiviral packaging plasmids (pRRE, pREV and pMD2.G) in 4:3:3:1 ratio in HEK293 Lenti-X cells for viral particle generation. 12h post-transfection, media change was given and incubated in general culture conditions for 72h. Viral particles were then collected by filtering the media with a 0.45 µM syringe filter. This viral particle filtrate was then added along with fresh media and Polybrene (5µg/mL) to HT1080 and MDA-MB-231 cells (for ind. hTERT construct) and HT1080, HCT116 WT cells and MDA-MB-231 cells (for ind. TRF2 variant construct); incubated for 24h. Post 24h, media containing viral particle was safely discarded and fresh media was added. Next, puromycin selection was given to the cells until the transduced population of cells could be stably maintained.

### Generation of TRF2 PTM variants using site-directed mutagenesis

The TRF2 post-translational modification variants were generated using site-directed mutagenesis by PCR method. The primers containing the required mutation (as documented in the table) were used to amplify the entire pCMV6 plasmid containing the TRF2-WT sequence. The PCR product was purified and transformed into DH511 E. coli cells. Then the mutated plasmid constructs were isolated and confirmed by Sanger sequencing.

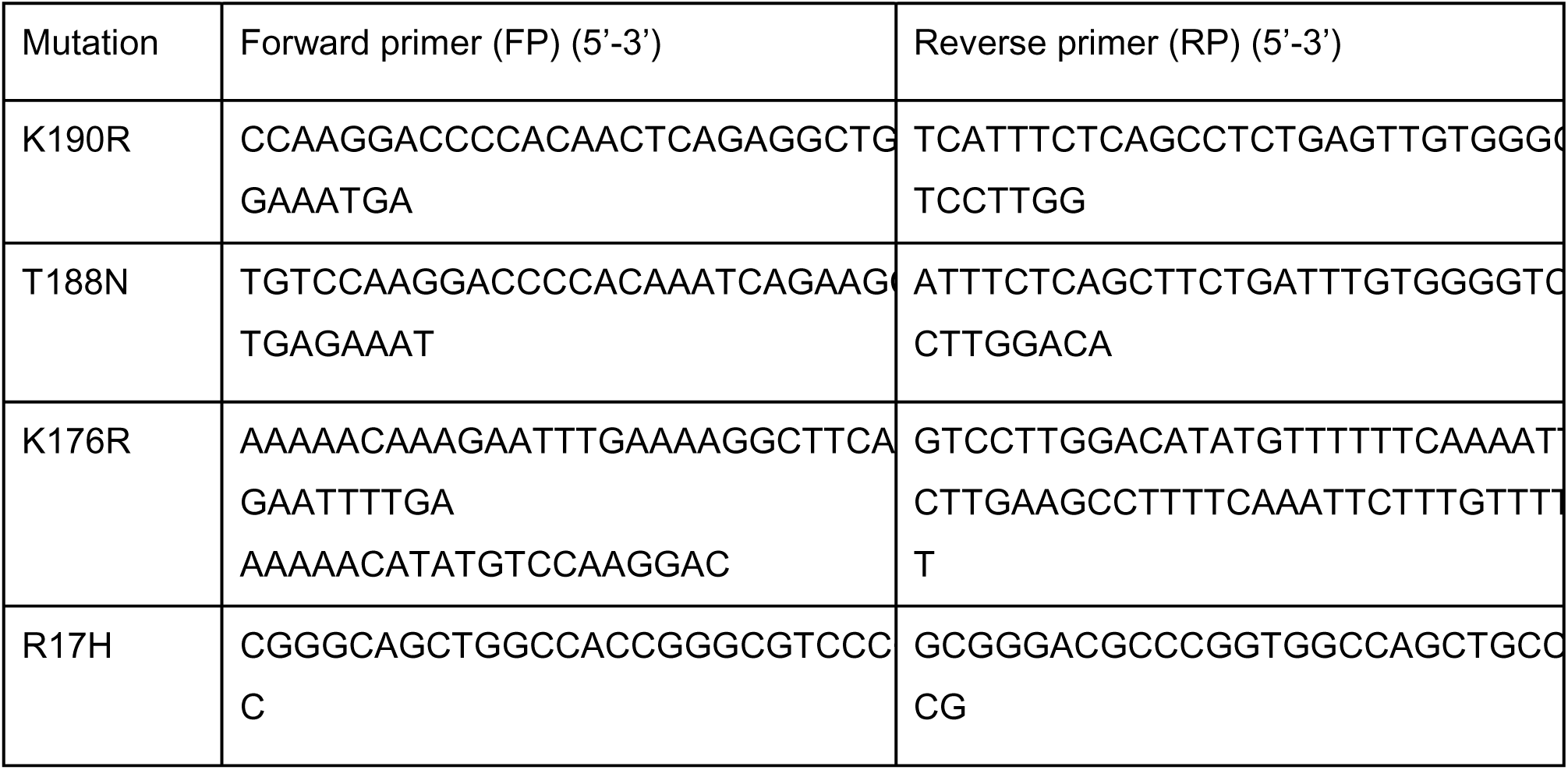

### Generation of xenograft tumour in NOD-SCID mice

Tumour xenograft generation in NOD SCID mice was outsourced to Vivo Bio Tech. Ltd – Telangana, India. 2.5 million cells of HT1080, and HT1080-LT each were injected subcutaneously in healthy male mice (4-6 weeks old) and monitored for tumour growth. Tumours were harvested by sacrificing mice when tumour growth reached an average volume of 600 mm^3^(± 100).

### Telomere length determination

Telomere length was determined using two methods - RT-PCR-based and Flow Cytometry-based. For flow cytometry-based evaluation using Agilent DAKO kit, 2 x10^6^ cells were divided into two fractions-for FITC labelled Telomeric PNA probe (Agilent DAKO kit) and unlabeled control cells. The cell fractions were washed and further labelled with PI for whole DNA content as per the manufacturer’s protocol. The fluorescence signal was acquired for at least 10,000 events on BD Accuri C6 flow cytometer. FCS files were then analyzed using FlowJo version 10.8.1 software. Fold change of telomere length relative to reference cell samples was calculated using Median Fluorescence signal intensity values of labelled and unlabeled sample preparations, according to the kit-provided formula.

For RT-PCR-based telomere length estimation, genomic DNA was isolated using the Wizard Genomic DNA Purification kit. At least 2 dilutions of genomic DNA in the range of 10ng-60ng were prepared for each sample and qRT-PCR was setup for telomere-specific primers and single copy gene 36B4 specific primer from O’Callaghan et al., 2008 and Cawthon et al 2002 respectively(72,73). Ct value of telomeric primer was compared to respective sample 36B4 and then fold change over the control sample was calculated.

The primers used are (5’ – 3’):

Tel F CGGTTTGTTTGGGTTTGGGTTTGGGTTTGGGTTTGGGTT
Tel R GGCTTGCCTTACCCTTACCCTTACCCTTACCCTTACCCT
36B4 F CAGCAAGTGGGAAGGTGTAATCC
36B4 R CCCATTCTATCATCAACGGGTACAA

### Telomerase activity using ELISA TRAP

ELISA TRAP was performed as described in Sharma et al 2021. Briefly, 1×10^6^ cells were lysed using mild lysis buffer CHAPS supplemented with RNase inhibitor to prevent loss of enzymatic activity (due to *hTERC* degradation) and PIC to prevent protein degradation. Protein concentration in the lysate was estimated and diluted to 0.5 µg/µL concentration. 2µg of protein was then used to set up telomerase repeat amplification protocol (TRAP) PCR(74). This was performed using a PCR master mix provided with ROCHE TeloTAGGG Telomerase PCR ELISA kit which uses 3’ biotinylated telomeric sequence repeat primers, P1-TS primers. Telomerase from the lysate extends the P1-TS primer which in the next step acts as a template for amplicon generation using P1-TS primer and (TTAGGG)6 repetitive sequences-P2. After this PCR, dioxygenin (dig) tagged telomere-specific detection probes are hybridized to denatured PCR products. These tagged biotinylated PCR products are then bound to streptavidin-coated plates and quantified using anti-dig-Peroxidase conjugated antibody, which produces coloured product upon metabolizing substrate, TMB).

### ChIP (chromatin immunoprecipitation)

Relevant Primary antibodies (TRF2 Novus Biologicals #NB110-57130, REST Sigma Aldrich #17-641, EZH2 Cell Signal Technology # 5246, H3K27me3 Abcam #ab6002, H3 Abcam # ab1791, FLAG Merck #F3165) and suitable IgG (from Millipore, for Isotype control or Mock) were used to perform the ChIP assays as per protocol reported in Mukherjee et al. (2018)(21). Briefly, 3 x 10^6^ cells were harvested and fixed with 1% formaldehyde for 10min. Post washing with ice cold PBS, the fixed cells were lysed in SDS based lysis buffer with 2X mammalian Protease Inhibitor Cocktail (mPIC) and subjected to chromatin fragmentation to an average size range of 200-300bp using Bioruptor (Diagenode). 10% fragmented chromatin was processed as Input fraction via Phenol-Chloroform-Isoamyl Alcohol (PCI) and Ethanol precipitation. Assay was performed with 3 µg of respective primary antibody incubated overnight on rotor at 4°C followed by immune complex pulldown with salmon sperm DNA saturated Protein G Magnetic Dyna Beads. Subsequent washes with low salt, high salt and then LiCl buffers were performed and beads resuspended in TE (Tris-EDTA pH 8) for proteinase K treatment at 55°C, 2h. Thereafter, PCI was used to phase - separate DNA in aqueous layer; separated fraction was incubated with equal volumes of isopropanol, 3M sodium acetate and glycogen overnight at -20°C. Pulldown DNA was then precipitated by centrifugation and pellet washed in 70% ethanol before air drying and resuspending in nuclease free water. Chromatin pulldown was validated by q-PCR analysis.

### Analysis of ChIP experiments

ChIP-q-PCR for TRF2, REST and EZH2 chromatin pulldowns was performed with equal amount of DNA for each fraction, determined by Qubit HS DNA kit. Then, fold change was calculated over IgG (Mock) by using average Ct values. In case of histone ChIP assays too, equal amount of DNA from each histone ChIP and its respective total H3 were used in qPCR. Ct values for Histone mark and total H3 were normalised to 1% Input and thereafter fold change over total H3 was calculated.

### Transfections

Cells were seeded a day before transfection at 70% confluency in a 6 well plate. 1.5µg plasmid per well was transfected by complexing with FUGENE at 1:3 (DNA: Reagent) ratio following manufacturer’s protocol.12h post transfection media change was given to the cells and later harvested for mRNA and protein analysis after 36h of media change.

### Real time PCR for mRNA expression

Total RNA isolation was performed using TRIzol Reagent (Invitrogen, Life Technologies) as per manufacturer’s instructions. RNA quantification and cDNA preparation was done using Applied Biosciences kit. Quantitative real-time PCR using SYBR Green based method (DSS TAKARA) was employed to estimate relative transcript levels of mRNAs with GAPDH as housekeeping control gene for internal normalization. Average fold change was calculated by difference in internally normalised threshold cycles (Ct) between test and control samples. All experiments were performed in three independent biological replicates; with technical triplicates of each primer pair and sample combination for q-PCR analysis.

In HT1080 ST/LT, Dox-HT1080 and Dox-MDA-MB-231 cells with overexpression of hTERT and *hTERC* (for HT1080-LT), primer pair specific for *hTERT* mRNA 3’UTR was implemented to capture the status of endogenous *hTERT* expression in addition to standard primers spanning exon 15/16 (full length) and exon 7/8 (active reverse transcriptase) forms. 3’UTR *hTERT* primers were designed with help from UTRdb 2.0.

mRNA primers (5’ – 3’):

hTERT 3’ UTR FP AATTTGGAGTGACCAAAGGTGT
hTERT 3’ UTR RP TATTTTACTCCCACAGCACCTC
hTERT exon 15/16 FP GGGTCACTCAGGACAGCCCAG
hTERT exon 15/16 RP GGGCGGGTGGCCATCAGT
hTERT exon 7/8 FP GCGTAGGAAGACGTCGAAGA
hTERT exon 7/8 RP ACAGTTCGTGGCTCACCTG
GAPDH FP TGCACCACCAACTGCTTAGC
GAPDH RP GGCATGGACTGTGGTCATGAG
TRF2 FP CAGTGTCTGTCGCGGATTGAA
TRF2 RP CATTGATAGCTGATTCCAGTGGT
18S FP TTCGGAACTGAGGCCATGAT
18S RP TTTCGCTCTGGTCCGTCTTG

### *hTERT* mRNA FISH

Untreated or transfected cells were seeded on coverslip for this experiment and processed according to Stellaris RNA FISH protocol. Briefly, cells were fixed with fixation buffer for 10min at RT. This was followed by PBS wash and permeabilization with 70% (v/v) ethanol. For Hybridization, 125nM of probe per 100ul of Hybridization buffer was mixed and added on to parafilm surface in a humidified chamber. Coverslips were then placed on this drop with cells side down (post wash in wash buffer A containing formamide), incubated in dark at 37°C overnight. After incubation, the coverslips were transferred to fresh 1ml Wash Buffer A for 30min. This was followed by nuclear staining with DAPI in the same buffer, in the dark for 30min maximum. Finally, the coverslip was washed in Wash Buffer B for 5min and then mounted on the slide with anti-fade mounting reagent.

Post slide processing, fluorescence signal for Quasar 670 *hTERT* RNA probe and DAPI were acquired on Leica SP8 confocal microscopy system. Implementing the default Rolling ball radius feature of open-source image analysis software Fiji ImageJ, background signal subtraction of each image frame was performed. Thereafter signal from n=30 cell nucleus (manually marked) was quantified.

### Immunofluorescence microscopy

Adherent cells were seeded on coverslips at a confluency of ∼ 70% before transfecting them with required TRF2 overexpression plasmids. At the harvesting time-point, cells were fixed using freshly prepared 4% Paraformaldehyde and incubated for 10 min at RT. Cells were permeabilised with 0.1% Triton™ X-100 (10 min at RT). The cells were treated with blocking solution (5% BSA in PBS + 0.1% Tween 20 (PBST)) for 1h at RT. All the above steps were followed by three washes with ice cold PBS for 5 min each. Post-blocking, cells were treated with anti-FLAG antibody (Merck #F3165,1:1000 diluted in 5% PBST) and incubated overnight at 4 C in a humid chamber. Post-incubation, cells were washed alternately with PBS three times for 5min and probed with secondary Ab mouse Alexa Fluor® 488 (A11059,1:3000) for 2 hrs at RT, in the dark. Cells were washed again with PBS three times and mounted with Prolong® Gold anti-fade reagent with DAPI. Images were taken on Leica TCS-SP8 confocal microscope.

### TRF2 protein purification

1.5 x 10^6^ HEK293T cells (∼60% confluency) were transfected with 3µg of TRF2-WT or TRF2-R17H overexpression pCMV6 plasmid. 24h post-transfection, cells were treated with G418 selection for 36-48h. Post this period of overexpression under selection pressure, the cells were harvested and resuspended in around 3-4mL ice-cold TBS (150mM NaCl, 50mM Tris-Cl, pH7.4) with 2X mammalian Protease Inhibitor Cocktail (mPIC). The cells were then sonicated to lyse open at 15s ON/ 30s OFF cycles for 5min (until turbidity was reduced), at 4°C. Cell debri was then separated by centrifuging at 13,000rpm, 5min, at 4°C. M8823 M2 FLAG Dynabeads were prepared for incubation with the cell lysate as follows: 40µL of the bead slurry was washed/ equilibrated twice in TBS by loading on 1.5mL tube magnetic stand, kept on ice. This equilibrated bead solution is then added to the cell lysate and incubated overnight at 4°C.

Post incubation of lysate with M2 FLAG bead for binding, the supernatant solution was removed and beads (bound to tube on magnetic stand) were washed based on step gradient purification. 2 washes with 1mL each of 150mM, 300mM, 500mM and 1M NaCl containing TBS were given to remove non-specific proteins bound to beads. All the wash fractions along with the supernatant and purified bead-bound TRF2 fractions were then analysed by running a 10% SDS-PAGE and staining with Coomassie Brilliant blue.

### Histone H3 methyl transferase activity quantification assay

1µg of purified histone H3 N-terminal peptide (procured as a component of Active Motif kit Catalog No. 56100) was incubated overnight (12-14h), at 37°C in the presence of 8µM (working concentration) S-adenosyl-L-methionine, 0.6µg purified PRC2 complex of proteins (cat no.SRP0381) and in the presence/absence of M2 FLAG bead-bound TRF2 protein (WT or R17H variants) in 25µL of a histone assay buffer (component of ab113454 kit). After that, the protocol of ab115072 –Histone H3 (tri-methyl K27) quantification kit was followed. Briefly, 25µL of supernatant of the reaction mix was pipetted, added to the anti-H3K27me3 antibody-coated strip wells with 25 µL of antibody buffer (provided with kit ab115072) and incubated at room temperature for 2h; the wells were covered properly by Parafilm M to prevent evaporation. Post incubation, the wells were washed thrice with kit-provided wash buffer. Diluted detection antibody (as directed by kit) was then added to the wells and incubated for 1h at room temperature on an orbital shaker at 100 rpm. Next, the wells were washed thrice after aspirating the antibody solution. The suggested amount of Color Developer solution was added to the wells and incubated in the dark at room temperature for 2-10 minutes. The colour development (blue) was monitored across the wells of sample and standard solution. Once the color in the standard well containing a higher concentration developed a medium blue colour, 50 μL of Stop Solution was added to all wells to stop the enzymatic reaction. Upon stopping the reaction, a yellow colour develops and the absorbance reading at 450nm was noted within 15minutes on a microplate reader. Fold change in trimethylation over reference condition is determined by the following formula:

Fold change of Tri-methylation = (Treated (tested) sample OD – blank OD) / (Untreated (control) sample OD – blank OD)

Blank in case of experiment was the reaction mix without PRC2 complex and TRF2 protein forms.

### Western blot analysis

Western blot analysis for checking the integrity of the purified bead-bound protein was also carried out. An equal volume of bead (5µL) was loaded for both TRF2-WT and TRF2-R17H on a 10% SDS-PAGE and transferred on polyvinylidene difluoride (PVDF) membranes (Biorad). Membranes were blocked with 5% skimmed milk and then probed suitably with primary antibodies-anti-TRF2 antibody (Novus Biological), and anti-FLAG antibody (Merck). Anti-rabbit and anti-mouse HRP conjugated secondary antibodies from CST respectively were used. Blots were developed using BioRad ClarityMax HRP chemiluminescence detection kit and images acquired on the LAS500 GE ChemiDoc system.

### Gaussia-luciferase assay

Secreted Gaussia luciferase signal in the CRISPR-modified short telomere and control cells (WT, -124G>A and -146G>A -ST/LT) was quantified using Promega Gaussia luciferase kit according to the manufacturer’s protocol. The luminescence signal (arbitrary unit, a.u.) was normalised to total protein for each condition.

### Quantification and statistical analysis

All experiments were performed with biological replicates, and data are presented as the mean ± standard deviation (SD) unless otherwise stated. Statistical significance was determined using appropriate tests based on experimental design.

For comparisons between groups where samples were derived independently (e.g., comparisons between each parental cell line and its corresponding short or long telomere-altered counterpart, or between control and TRF2-overexpressing conditions), an unpaired two-tailed Student’s t-test was used. For experiments in which the same set of biological replicates was subjected to different treatments or time points (e.g., doxycycline induction and post-withdrawal), a paired two-tailed Student’s t-test was applied.

For comparisons across more than two time points or conditions (e.g., telomere length analysis or expression dynamics during doxycycline treatment), one-way ANOVA followed by Tukey’s Honestly Significant Difference (HSD) post-hoc test was used to identify statistically significant differences between groups.

For experiments involving two independent variables (e.g., Guassia luciferase assay across multiple conditions and time points, or in vitro histone methyltransferase activity under different treatments), two-way ANOVA was performed to assess the interaction effects. Where applicable, post-hoc comparisons were conducted to identify significant pairwise differences.

Statistical significance is indicated as follows: *p < 0.05; **p < 0.01; ***p < 0.005; ****p < 0.0001. All statistical analyses and data visualization were performed using GraphPad Prism version 8.0.2.

## Acknowledgments

The authors thank Balam Singh for management of laboratory and resources. We are thankful to all members of the S.C. laboratory for their suggestions/inputs and the IGIB Core Imaging Facility for its service facility. We acknowledge assistance from Dr. Arnab Mukhopadhyay and his lab members (National Institute of Immunology) for their cooperation. Research fellowships to ASG, SVM, DS, RD, JJ, SD, AKM, SSR, PP, SS, AKB, DK, AS, SS, SB (CSIR) and M.Y.(UGC) are acknowledged.

## Author contributions

Conceptualization, SC; Methodology, ASG, SVM, DS, RD, JJ, SD, AKM, MY, SSR, SS, AS, SS, RKM; Validation ASG, SVM, SB, AS, SD; Formal Analysis ASG; Investigation, ASG, SVM, DS, RD, JJ, SD, AKM, MY, PP, AKB; Resources ASG, AS, SS, RKM, AS, DK, SSR, SB; Data Curation, ASG; Writing – Original Draft Preparation, ASG; Writing – Review & Editing, SC and ASG; Visualization, ASG; Supervision, SC; Project Administration, ASG, SC; Funding Acquisition SC

## Data and materials availability

All data are available in the main text or the supplementary materials.

