## Supplemental figures for "Telomeres control human telomerase (*hTERT*) expression through non-telomeric TRF2"

**Supplementary Figure 1**

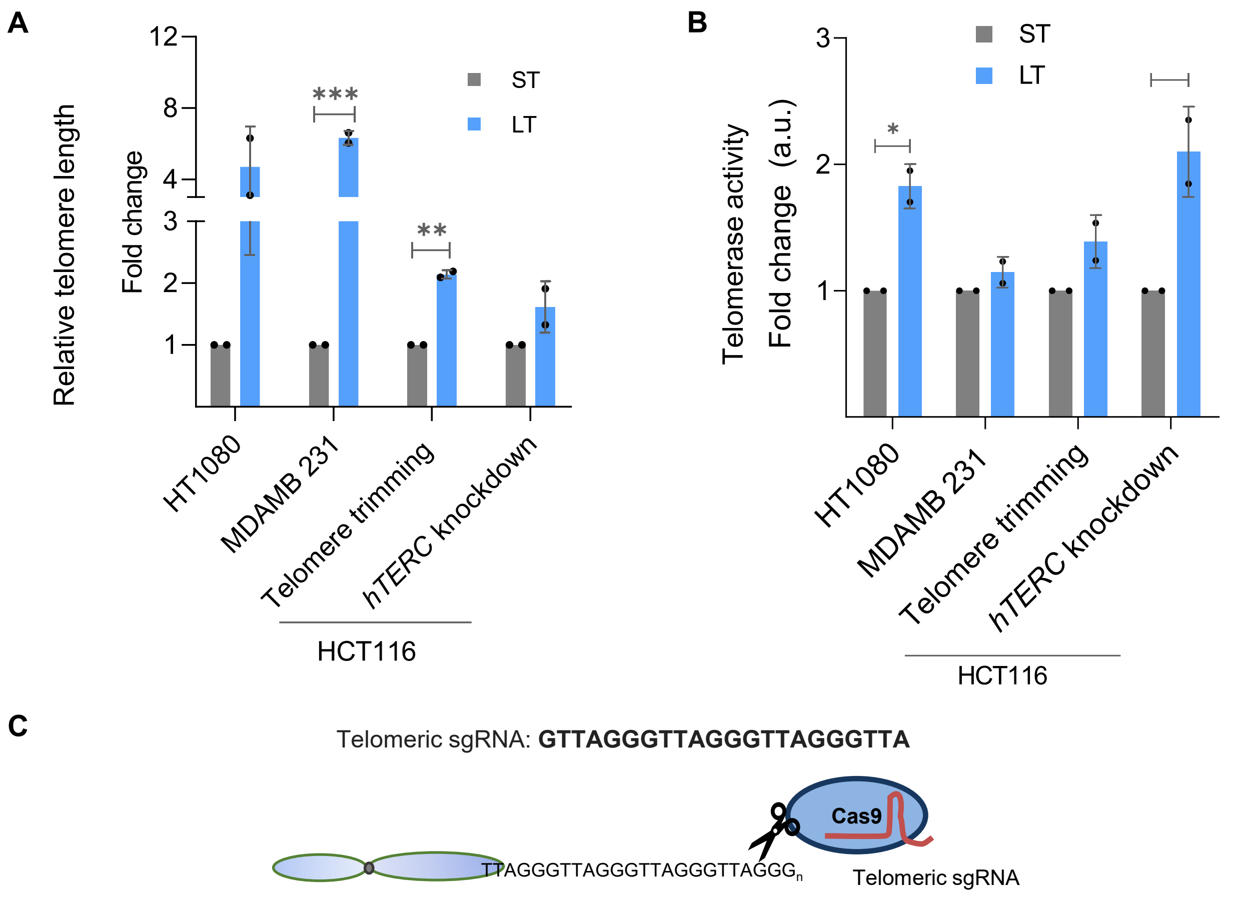


Supplementary Figure 1:

(A) Telomere length in isogenic cancer cell lines with short telomeres (ST, in grey) or long telomeres (LT, in blue) namely, HT1080- ST/LT, MDA-MB-231-ST/LT, and HCT116-ST/LT (Telomere trimming-Cas9 and *hTERC* knockdown) as determined by qRT-PCR based assay (See Methods).

(B) Telomerase activity in isogenic cancer cells with short telomeres (ST) or long telomeres (LT) namely, HT1080- ST/LT, MDAMB 231-ST/LT, HCT116-ST/LT and HCT116 p53 null -ST/LT, determined using telomerase-repeat-amplification-protocol (TRAP) followed by ELISA (see Methods);

(C) Scheme showing the telomeric specific sgRNA used to generate short telomere versions in HCT116, HEK293T CCR5 hTERT promoter insert cells and iPSCs using transient expression of Cas9- telomere targeting sgRNA plasmid.

Error bars represent ± SDs from the mean from two independent biological replicates. p values are calculated by unpaired t-test. (*$p < 0.05$, **$p < 0.01$, ***$p < 0.005$, ****$p < 0.0001$).

### Supplementary Figure 2

**A**

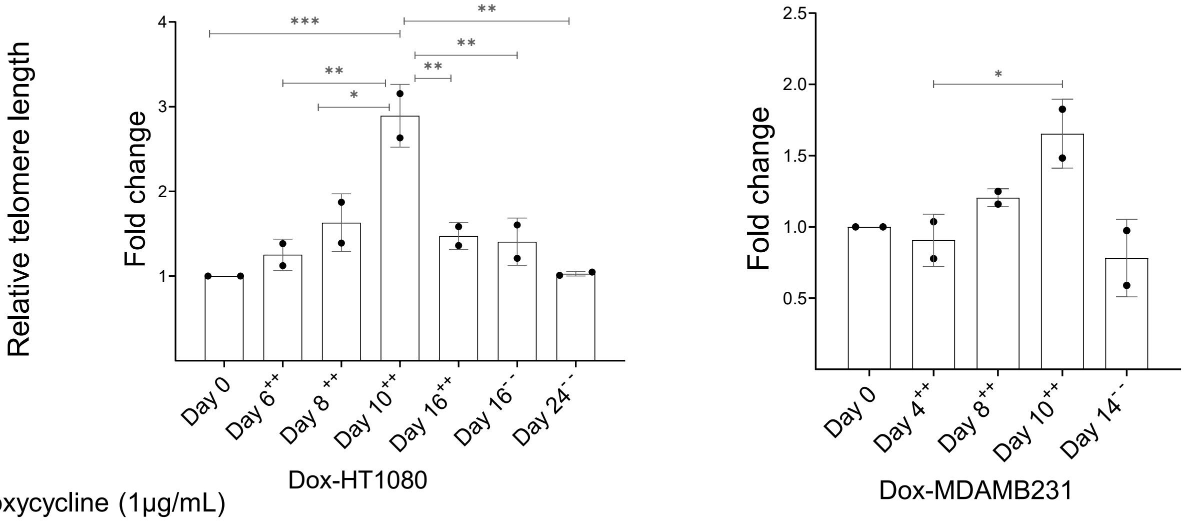


Doxycycline (1µg/mL)

++ Present

- -  Absent

**B**

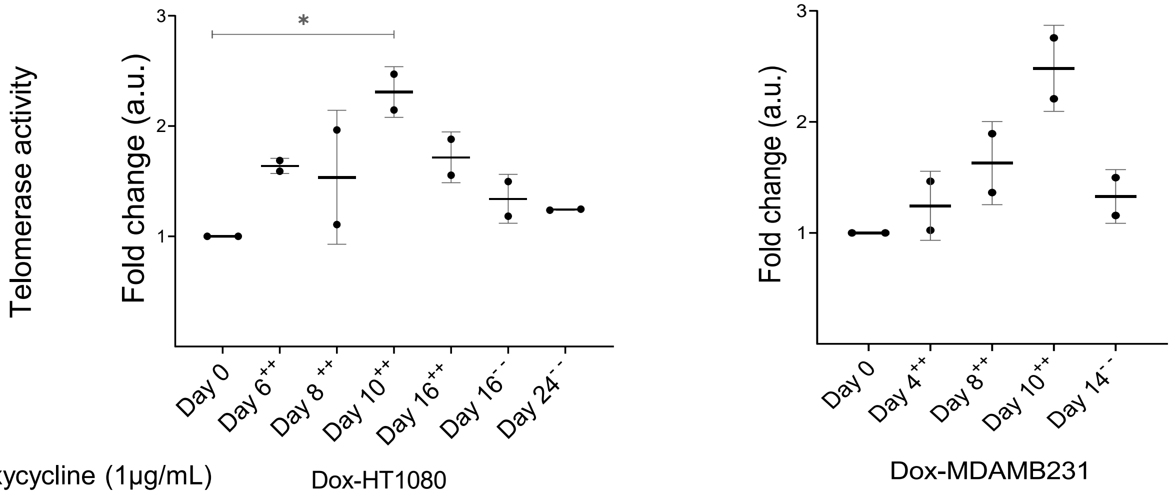


Doxycycline (1µg/mL)

++ Present

- -  Absent

**Supplementary Figure 2:**

(A) Relative fold change in telomere length at multiple day intervals (as indicated) in Dox-HT1080 cells (Left panel) and Dox-MDAMB231 (Right panel) as determined by qPCR- based telomere length detection method Telomeric signal normalised over single copy gene, 36B4 for qPCR-based analysis. ++/-- denotes the presence/ absence of dox at the indicated day points.

(B) Telomerase activity at multiple-day intervals (as indicated) in Dox-HT1080 cells (Left panel) and Dox-MDAMB231 (Right panel) cells determined using telomerase-repeat-amplification-protocol (TRAP) followed by ELISA (see Methods); ++/-- denotes the presence/ absence of dox at the indicated day points.

All error bars represent ± SDs from the mean from two independent biological replicates. One-way ANOVA followed by post-hoc tests (Tukey's HSD) was performed to compare means across time points in figures (A,B). (*$p < 0.05$, **$p < 0.01$, ***$p < 0.005$, ****$p < 0.0001$).

**Supplementary Figure 3**

A

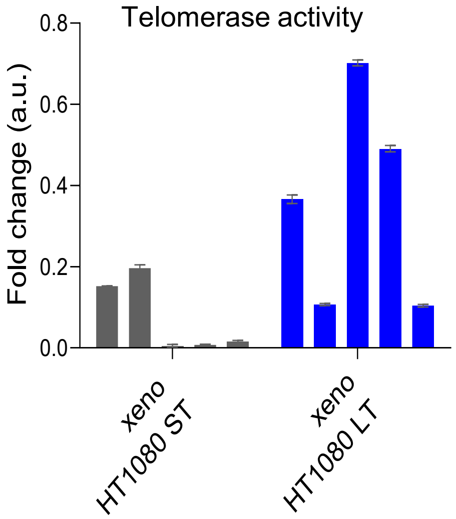


**Supplementary figure 3:**

(A) Telomerase activity in xenograft tissues of HT1080-ST and HT1080-LT cells

**Supplementary Figure 4**

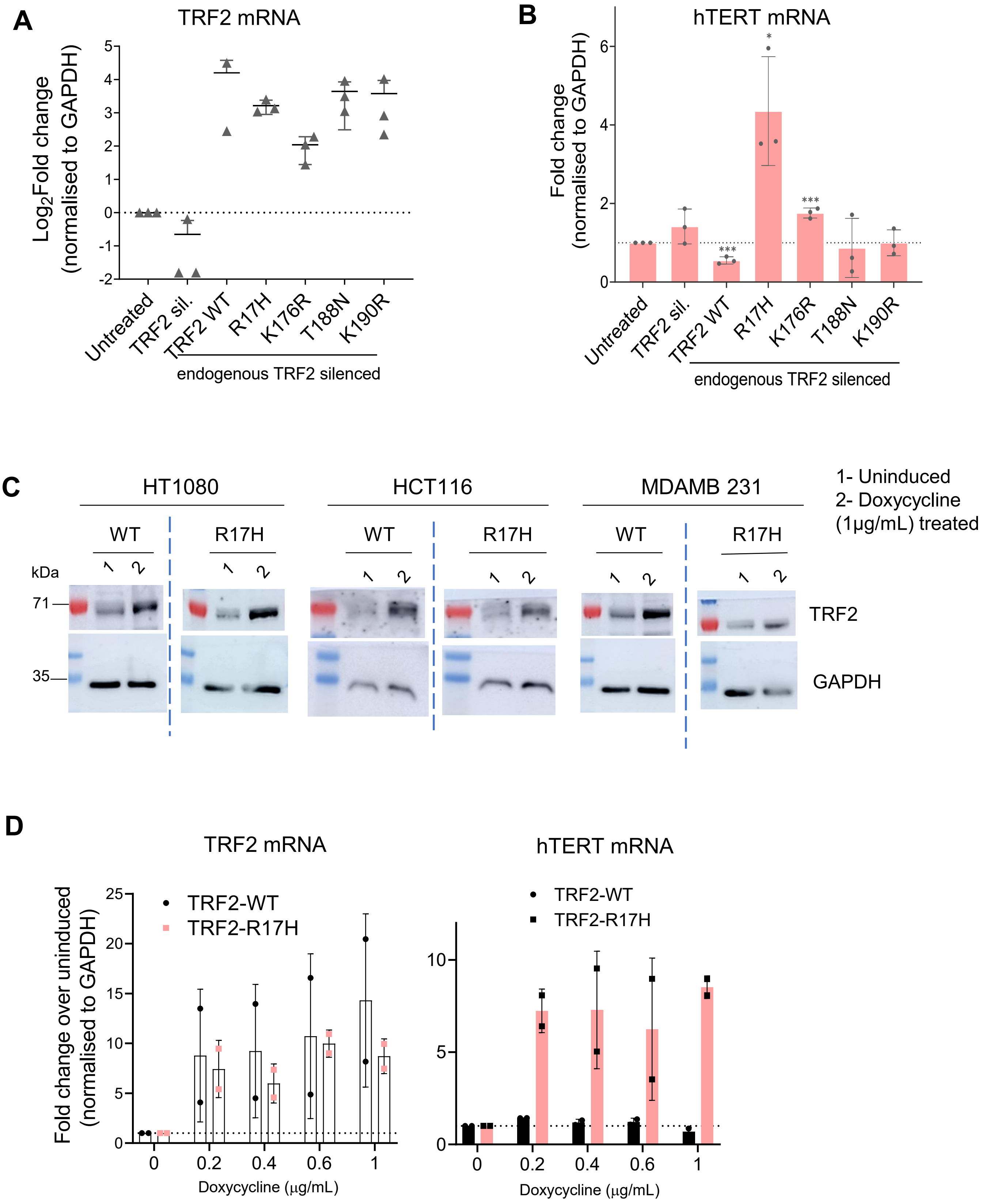

#### Supplementary Figure 4

**E**

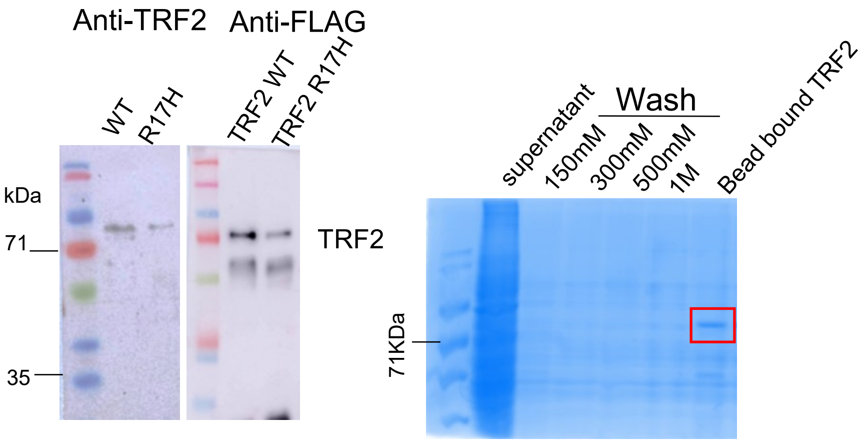


**F**

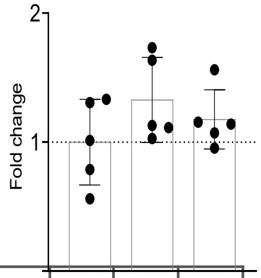


| | 1 | 2 | 3 |
|---|---|---|---|
| Histone H3 | + | + | + |
| S-Adenosyl Methionine | + | + | + |
| PRC2 protein complex | - | + | + |
| Empty FLAG bead | - | - | + |

**Supplementary Figure 5:**

(A) TRF2 mRNA levels in HT1080 cells as TRF2 PTM variants are overexpressed under endogenous TRF2 silenced conditions using shRNA

(B) hTERT full-length mRNA transcript (exon 15/16) levels by RT-PCR in HT1080 cells upon overexpression of TRF2 PTM variants as depicted in (A) under endogenous TRF2 knockdown condition (B).

(C) TRF2 protein induction with Dox treatment in HT1080, HCT116 and MDAMB 231 TRF2 inducible lentiviral stable cells, confirmed by Western blot analysis (Mol. Wt. ladder used in HT1080 and MDAMB 231 is Puregene 4 color Prestained Protein Ladder,10-180 kDa and that of HCT116 is G Biosciences PAGEmark Tricolor PLUS.

(D) Dose-dependent Dox induction of TRF2-WT and R17H variant (Left graph) in stable inducible TRF2 HT1080 cells to check hTERT full-length mRNA transcript (exon 15/16) levels by RT-PCR (right graph).

(E) Purified TRF2 WT and TRF2 R17H protein from HEK 293T cells as developed by anti-TRF2 and anti-FLAG antibodies (left panel) and representative Coomassie Brilliant Blue (CBB) gel for protein purification protocol. The lower band in the anti-FLAG blot is of bead-bound FLAG peptide.

(F) H3K27 trimethylation levels in in vitro histone methyl transferase assay with empty (TRF2 unbound) FLAG beads.

Error bars represent ± SDs from the mean from two independent biological replicates. p values are calculated by unpaired t-test in 5B. (*p $< 0.05$, **p $< 0.01$, ***p $< 0.005$, ****p $< 0.0001$).
